# Recombination of standing variation in a multi-hybrid swarm drove adaptive radiation in a fungal pathogen and gave rise to two pandemic plant diseases

**DOI:** 10.1101/2021.11.24.469688

**Authors:** Mostafa Rahnama, Bradford Condon, Joao P. Ascari, Julian R. Dupuis, Emerson Del Ponte, Kerry F. Pedley, Sebastián Martinez, Barbara Valent, Mark L. Farman

**Affiliations:** Department of Plant Pathology, University of Kentucky, Lexington, KY, 40546 USA; Departamento de Fitopatologia, Universidade Federal de Viçosa, Campus Universitário s/n 36570-900, Viçosa-MG, Brazil; Department of Entomology, University of Kentucky, Lexington, KY, 40546 USA; USDA/ARS/Foreign Disease Weed Science Research Unit, 1301 Ditto Ave. Bldg. 1301, Fort Detrick, MD 21702; Instituto Nacional de Investigación Agropecuaria (INIA), Laboratorio de Patología Vegetal, INIA Treinta y Tres, Ruta 8, Km 281, 33000 Treinta y Tres, Uruguay; Department of Plant Pathology, Kansas State University, Manhattan, KS, 66506, USA

## Abstract

Adaptive radiations fuel speciation and are characterized by rapid genetic diversification and expansion into new ecological niches. Historically, these processes were believed to be driven by selection on novel mutations but genomic analyses now indicate that standing variation and gene flow often have prominent roles. How “old” variation is combined, however, and its resulting genetic architecture within newly-adapted populations is not well understood. We reconstructed a recent radiation in the fungus, *Pyricularia oryzae,* that spawned a population pathogenic to eleven grass genera, and caused two new plant diseases: wheat blast - already a serious threat to global agriculture - and gray leaf spot of ryegrasses. We show that the new population evolved in a multi-hybrid swarm using only the standing variation that was present in seven individuals from five distinct, host-specialized lineages. Sexual and parasexual recombination within the swarm reassorted key host-specificity factors and generated more diversity in possibly just a few weeks than existing lineages had accumulated over hundreds to thousands of years. We suggest that the process was initiated by sexual opportunity arising when a fertile fungal strain was imported into Brazil on *Urochloa* introduced as forage for beef production; and we further contend that the host range expansion was largely fortuitous, with host selection playing little, if any, role in driving the process. Finally, we believe that our findings point to an overlooked role for happenstance in creating situations that allow organisms to skirt rules that would normally hold evolution in check.

**Commercial Endorsement Disclaimer:** Mention of trade names or commercial products in this publication is solely for the purpose of providing specific information and does not imply recommendation or endorsement by the U.S. Department of Agriculture.

**Equal Opportunity/Non-Discrimination Statement:** USDA is an equal opportunity provider and employer.

## INTRODUCTION

Adaptive radiations fuel speciation and are characterized by rapid genetic diversification and expansion into new ecological niches. Early evolutionary theory predicted such events are driven primarily by new mutations, with gene flow being counterproductive (*1, 2*). However, many radiations have occurred over timeframes far too short for the incremental accumulation of beneficial variants; and genomic studies have found that alleles associated with adaptation and speciation often vastly predate the radiation events themselves (*3, 4*). This, along with evidence of allelic re-assortment via admixture (*5, 6*), suggests that standing genetic variation and gene flow may serve more prominent roles (*7, 8*), but how “admixture variation” arises, and its genetic architecture within newly-adapted populations, is not well understood. Two recently emerged plant diseases caused by the fungus *Pyricularia oryzae* (syn. *Magnaporthe oryzae*) bear hallmarks of having arisen as part of an adaptive radiation. The short time since disease emergence, combined with a relatively small, haploid genome, and a wealth of information on the global pathogen population, offered an excellent opportunity to reconstruct the causal pathogen’s evolution.

First reported in 1985 in the Paraná State of Brazil (*9*), wheat blast (WB) (Fig. 1A) can cause 100% crop loss in some years. The disease quickly spread to surrounding countries (*10–12*) and, until very recently, was restricted to South America. However, major outbreaks in Asia in 2016 and 2017 (*13, 14*), and the pathogen’s arrival in Africa (*15*), have now elevated WB as a major concern for global agriculture. Shortly after WB’s emergence, another new disease caused by *P. oryzae* was reported on perennial ryegrass (*Lolium perenne*) turf used in Pennsylvania golf course fairways (*16*). Known as gray leaf spot (GLS) (Fig. 1B), the disease went on to cause major epidemics in the mid-to late 1990s, affecting perennial ryegrass and tall fescue (*Lolium arundinaceum*) throughout the central U.S. (*17*). The first reports of *P. oryzae* on *Lolium* grasses, however, were for isolated outbreaks that occurred in late 1971 on annual ryegrass (*Lolium multiflorum*) forage in Mississippi and Louisiana (*18, 19*). Interestingly, the fungal populations causing these two new diseases are closely related to one another, yet quite distinct from other well-established populations, which led to the proposition that they are a new species (*20*). However, evidence of gene flow has raised this status into question (*21, 22*).

**Figure 1.**
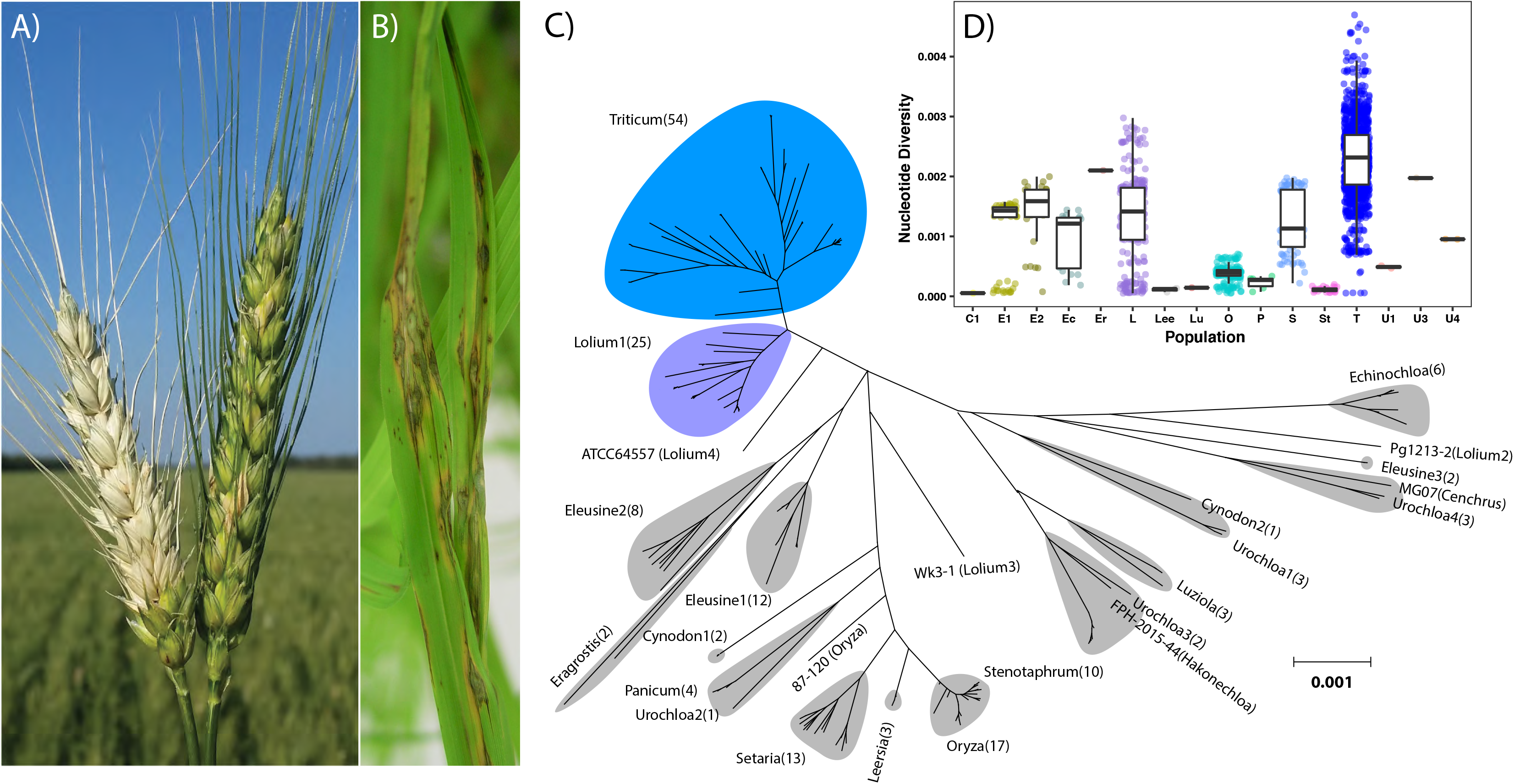
Phylogenetic relationships between *P. oryzae* host-specialized populations. A) blasted wheat head (left) and healthy head (right). B) gray leaf spot on tall fescue (*Lolium arundinaceum*). C) Phylogenetic tree showing pairwise distances between isolates. Phylogenetic clades (= lineages) are labeled according to the host of isolation, with numerical suffixes indicating order of discovery. Numbers of strains in each lineage are indicated in parentheses. Lineage memberships are provided in Table S1. Gray shaded areas show how phylogenetic lineages grouped according to DAPC (see below) (Fig. S3). D) Boxplot showing within-population, pairwise nucleotide diversity (∏), with mean and quartile boundaries.

*P. oryzae* is best known as a global pathogen of rice but also infects other crops such as oats, barley, millets, as well as a range of turf and weedy grasses (*23*). Normally, however, it exhibits strong host specificity and, although cross-infection of multiple hosts can been demonstrated using inoculation assays (*24*), the phylogenetic clustering of isolates by host of origin ((*21, 25*); Figure 1C) indicates that cross-infection is rare in nature. Curiously, this is not true for the phylogenetic lineages causing WB (PoT – *P. oryzae Triticum*-adapted lineage) and GLS (PoL-*P. oryzae Lolium*) because member isolates have also been recovered from oats and a number of weedy grasses (*21, 26–28*). Thus, the events underlying WB and GLS emergence spawned adaptations that furnished the pathogen with a dramatically enhanced host range.

Current understanding of WB evolution centers on the discovery that related strains from oats and *Lolium* species - are unable to infect wheat because they possess the *PWT3* host-specificity gene that confers avirulence towards wheat cultivars carrying *Rwt3* resistance (*29*). To explain the presence of functional and non-functional PWT3 alleles within the WB population, it was proposed that widespread cultivation of Anuhuac - a wheat cultivar lacking *Rwt3* - in the early 1980s supported an initial host jump from *Lolium* onto wheat, with nearby resistance cultivars then imposing selection for the eventual loss of *PWT3* (*29*). However, application of distance-based phylogenetic methods to characterize the populations revealed that PoT adaptation to wheat was accompanied by explosive genetic divergence - as was the adaptation of PoL to *Lolium* (*21*) (this study). A simple host jump through the loss of a single host specificity gene is not expected to generate such extreme genetic diversity, nor a general relaxation of host specificity and, therefore, these phenomena pointed to a much more complex evolutionary scenario.

## RESULTS

### The wheat blast and gray leaf spot pathogens evolved in an adaptive radiation that generated fungal populations with exceptionally wide host ranges

An updated phylogenetic tree built using genome-wide, pairwise distances showed that branch lengths within the PoT and PoL lineages were much longer than expected for populations that we predicted to have evolved shortly before gray leaf spot and wheat blast first emerged in 1971 and 1985, respectively (Figure 1C). Consistent with the tree, the PoT and PoL lineages showed far greater nucleotide diversity than was present in most of the other host-specialized lineages (Fig. 1D). For comparison, the rice (*Oryza*)- and foxtail (*Setaria*)-infecting lineages have existed for hundreds and possibly thousands of years, respectively (*30, 31*). Along with their enhanced genetic diversity, the PoT/PoL lineages showed exceptionally broad infection capabilities with member isolates being recovered from nine other grass genera, including oats (*Avena*), cheatgrass (*Bromus),* buffelgrass *(Cenchrus)*, crabgrass (*Digitaria),* barnyardgrass *(Echinochloa),* goosegrass (*Eleusine)*, balsamscale grass (*Elionurus*), Natal grass (*Melonis/Rynchelytrum*), and signalgrass (*Urochloa)* (*20*) (Fig. S1). Thus, the emergence of wheat blast and gray leaf spot appear to have been part of a broader adaptive episode that was driven by extremely rapid genome diversification. Accordingly, we reasoned that the key steps in WB/GLS evolution could be elucidated by using phylogenomic approaches to identify the processes driving such rapid sequence divergence. Another key point that arises from the expanded phylogenetic tree - one that speaks to implications of the new population’s broad host range, is that several host specificities have evolved many times independently in *P. oryzae*, including to *Cynodon* (2 lineages), *Eleusine* (3 lineages), *Lolium* (4 lineages), and *Urochloa* (4 lineages) (Figure 1C).

### The PoT and PoL lineages contain multiple alleles/variants for several marker loci that were inherited from at least five other host-specialized *P. oryzae* lineages

A first clue to the rapid diversification in PoT and PoL came from an analysis of haplotype divergence along the chromosomes. Pairwise comparisons between isolates from the “low divergence” rice blast population revealed that variation was distributed evenly along the chromosomes (Figure 2A, upper panel), with an average haplotype divergence (AHD) of 0.0127 across windows of 2,000 SNPs. The spikes are due to short, non-PoO introgressions in individual strains (e.g. the white arrowhead shows a spike caused by a PoU1 introgression in strain TH3; and the black one, a PoSt introgression in PH0014-rn). In contrast, alignments between PoT/PoL members showed extended regions of complete haplotype identity for most pairwise comparisons (AHD for windows 375 to 625 = 0.00051) -reflective of their recent origins - and these were interspersed with highly divergent regions. Critically, however, not all comparisons showed high divergence in these regions, as indicated by an abundance of plot lines tracking the x-axis (Figure 2A, lower panel).

**Figure 2.**
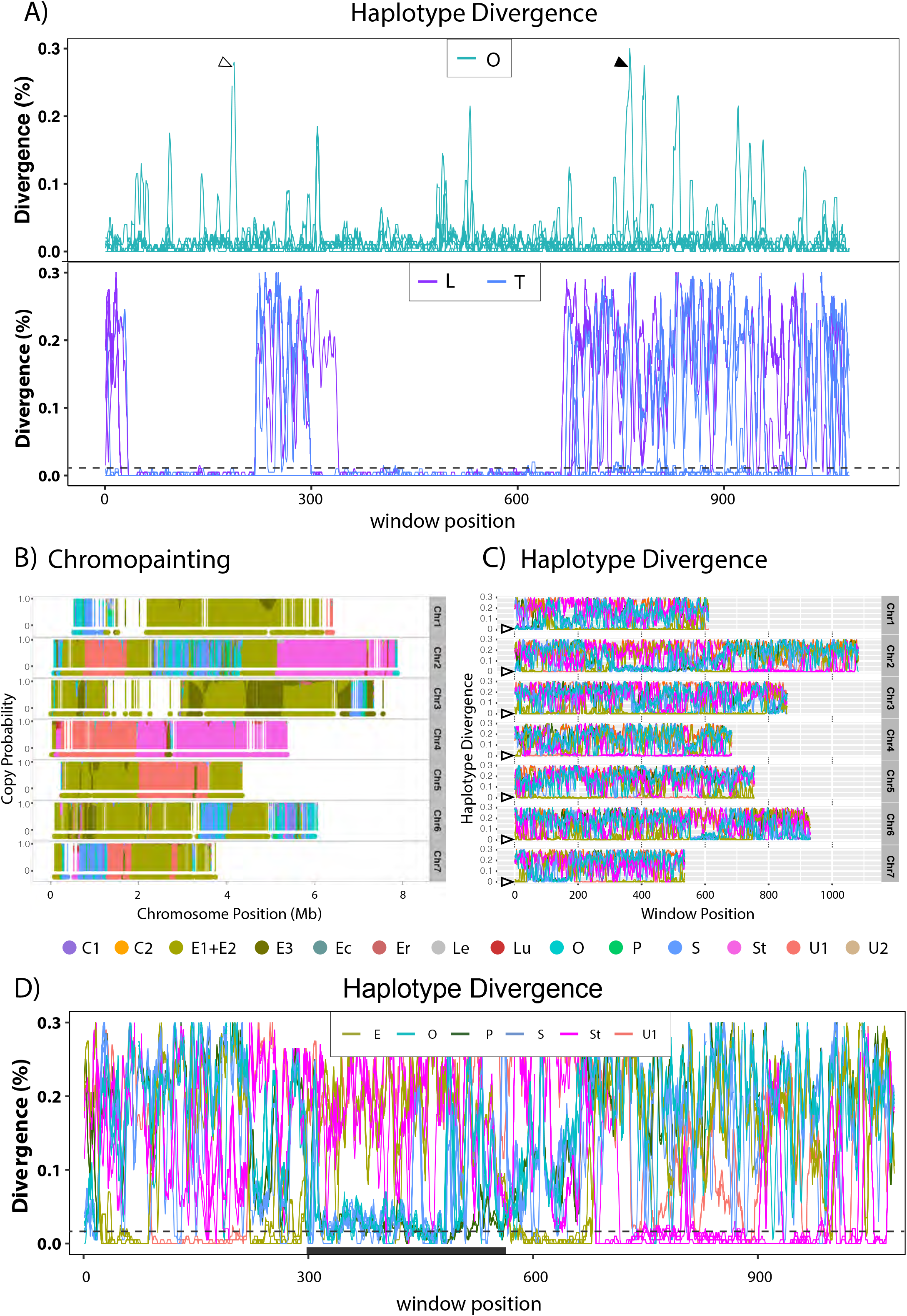
The genome of the wheat blast reference strain is a mosaic of sequences inherited from five host-specialized populations. A) Top panel: Chromosome 2 haplotype divergence between rice pathogen, Guy11, and other rice blast isolates (n = 15). Plots are superimposed to capture total variation within the PoO lineage. Bottom panel: Chromosome 2 haplotype divergence between B71 and PoL (n = 23) and PoT (n = 52) isolates on chromosome 2. Highly divergent regions in the chromosome are interspersed with regions showing almost zero divergence. B) Chromopaintings of B71 chromosomes with stacked plots showing copy probabilities for all candidate donors as listed in the legend below. Condensed plots at the bottom show most probable donor for SNPs with high confidence ancestry calls. Note: chromosome segments inherited from PoX show up as alternating blocks of blue/green shades because, without sampling a member of the actual PoX donor population, these segments are variously inferred as having ancestry to closely related populations: PoO/PoS/PoLe and PoP. C) Haplotype divergence between B71 and comparator strains from suspected donor lineages. A line is plotted for each B71 x test strain comparison, with line colors showing the lineage affiliations of the comparators. Note the extremely low haplotype divergence relative to the donor lineages inferred by the chromopaintings, and significantly greater divergence relative to other lineages. D) Haplotype divergence analysis identifies contributions from a “cryptic” population. Note that in most windows, the B71 chromosome segments inherited from the PoE, PoSt and PoU1 donors, B71, show zero haplotype divergence relative to at least one donor lineage member and divergence relative to other members is generally below dotted line. In contrast, the regions marked with the black bar shows the highest similarity (but not identity) to members of PoO, PoP, and PoS, yet significant divergence relative to the next most closely related lineages.

In seeking to explain the regions of exceptional nucleotide divergence, we recalled prior work showing that the PoT and PoL lineages possess up to four alleles/variants for several phylogenetic markers (*21*); and because nearly all of the variants were found in other host-specialized populations, this pointed to the introgression of DNA from several sources. To confirm this suspicion, we built phylogenetic trees using 20 kb regions surrounding CH7BAC7 and MPG1. This revealed that PoT/PoL actually had four variants at each locus. Two of the CH7BAC7 variants found in PoT/PoL showed perfect sequence identity with variants present in members of divergent populations infecting signalgrass (lineages PoU1 and PoSt). The third was a variant of the PoSt sequence, while the fourth was most closely related to, but slightly different from, sequences from rice (PoO) and foxtail (PoS) pathogens (fig. S2A). Similarly, three of the MPG1 variants shared perfect sequence identity with sequences found in isolates from goosegrass (PoE1) and signalgrass (PoU1 and PoSt), and the fourth, again, was most similar to, but different from, those found in PoO/PoS (fig. S2B). From these data, we concluded that the PoT and PoL lineages likely contain sequences introgressed from at least five pre-existing *P. oryzae* lineages: PoE1, PoSt, PoU1, PoU3, and a lineage related to PoO/PoSt. Furthermore, despite the fact that CH7BAC7 and MPG1 reside just 1 Mb apart on chromosome 7, the trees showed many incongruencies (fig. S2), pointing to recombination between the two loci.

### The wheat blast genome was assembled entirely from sequences inherited from pre-existing host-specialized lineages with former histories of divergent evolution

Extrapolating from the above findings, we surmised that the PoT and PoL genomes might comprise chromosome segments donated entirely by other host-specialized forms of the fungus. To explore this possibility, we first used discriminant analysis of principle components (DAPC) (*32*) to assign 96 non-PoT/PoL isolates (Table S1) to 16 discrete lineages (Fig. S3). Next, we used a chromosome-level reference assembly for PoT isolate B71 to call SNPs for the 96 candidate donors and employed ChromoPainter to paint the B71 chromosomes according to the likelihood that a given segment was contributed by a given donor lineage. This revealed the entire B71 genome to be a mosaic of sequences donated by pre-existing lineages, with the primary donors being PoE1/2, PoO, PoS, PoSt and PoU1 (Figure 2B).

To confirm the ChromoPainter results, we used the custom-built program, ShinyHaplotypes, to analyze patterns of haplotype divergence along the chromosomes when comparing the B71 genome against all possible donor isolates. This revealed near perfect haplotype identity between B71 and isolates from the donor populations predicted by ChromoPainter, and significant divergence relative to the next closest lineage(s) (Figures 2C). The only exceptions were haplotype blocks that ChromoPainter predicted to have been donated by the PoO/PoS lineages. These chromosome segments were clearly most similar to PoO and PoS but they exhibited greater divergence from B71 than the segments donated by PoE1, PoSt and PoU1 (Figure 2D) and, thereby, reflected the pattern seen with the PoO/PoS-like alleles in the CH7BAC7/MPG1 phylogenetic trees (Fig. S2). Considering that all the other “admixture” blocks showed near perfect identity to their predicted donors, and even the PoO/PoS-like segments showed almost perfect sequence identity when comparing different PoT/PoL strains (compare Figure 2A with C & D), this pattern is most economically explained by contributions from a “cryptic” population (“PoX”) that is phylogenetically related to PoO/PoS and infects a host that has not yet been sampled.

### PoT/PoL evolution was driven by re-assortment and recombination of chromosome segments in a multi-hybrid swarm with donations from five host-specialized populations

Having established that the B71 genome is a mosaic of donated chromosome segments, we surmised that this is likely true for the other PoT/PoL isolates. Furthermore, to explain the variable nucleotide divergence patterns (Figure 2A, bottom panel), the multiple alleles found for many marker loci, and the different ancestries of linked markers (Fig. S2), we hypothesized that PoT/PoL essentially constitutes a multi-hybrid swarm, where multiple admixture events created a genetic pool that contained significant proportions of the various donor genomes, and the constituent chromosomes were then differentially assorted/recombined through matings among swarm members. A neighbor-net phylogenetic network constructed using SplitsTree (*33*) showed many reticulations within the PoL and PoT clades, consistent with extensive recombination among constituent isolates (Fig. S4). Even more compelling evidence of recombination, however, came from chromopainted genomes for representative isolates from each of 43 distinct chromosomal haplotypes (11 for PoL and 31 for PoT, Table S2), which showed that each haplotype comprises variable combinations of chromosome segments inherited from each of the previously-identified donor populations (Figure 3). Also revealed was a significant contribution of chromosome 4 DNA from a new population found on watergrass (*Luziola spp.,* PoLu) and, although the Chromopainting makes it appear that PoLu segments are interspersed with PoSt DNA, ShinyHaplotypes shows that PoLu is another hybrid population whose members contain variable blocks of sequences with PoSt and PoU1 heritage, along with a third unknown contribution (Fig. S5). Thus, the alternating PoLu/PoSt segments on chromosome 4 likely constitute a single donation from just one PoLu isolate that is, itself, a hybrid. A similar situation involves PoE1 segments interspersed with short tracts with predicted PoE2/PoE3 heritage. Again, ShinyHaplotypes revealed most PoE1 isolates to contain introgressions of PoE2/3 DNA and, because the segments in question were present in nearly all isolates possessing the encompassing PoE1 DNA, this again pointed to the donor as being a single, recombinant PoE1 individual - one possessing PoE2/3 introgressions.

**Figure 3.**
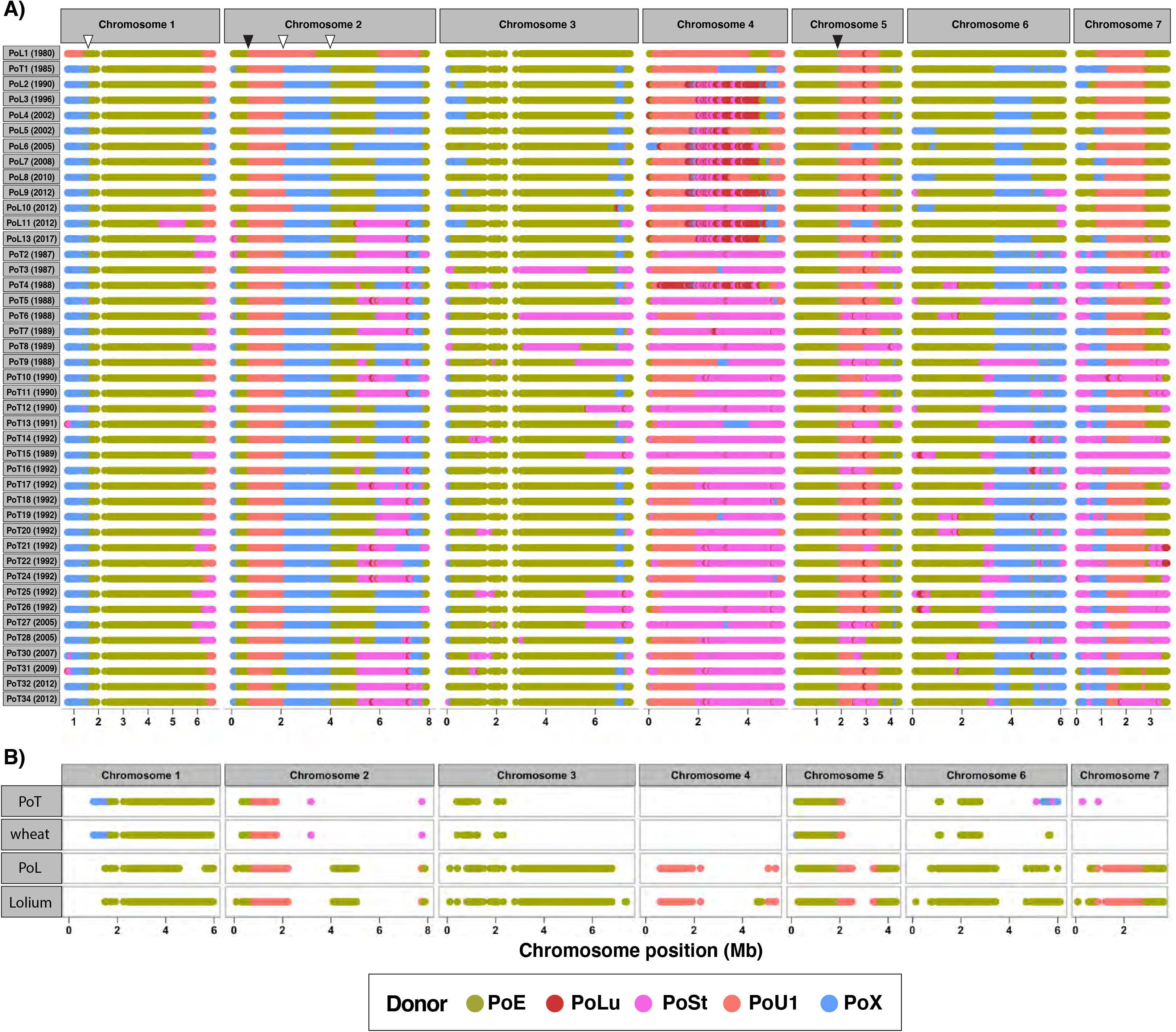
Chromosome paintings showing PoT/PoL haplotypes defined to date. A) For clarity, data points for populations E1, E2 and E3 were merged and segments predicted to come from PoEc, PoO, PoP or PoS were reassigned to PoX. Only contributions from the five main donor lineages are shown because predicted donations from other lineages were minor (< 1%), rare, and/or low confidence (data not shown). Donors are shown in the legend at the bottom. Two haplotypes were omitted due to poor assembly quality. B) Invariant chromosome segments in PoT and PoL. To illustrate how highly variable genome heritage has little impact on affinity for the wheat and *Lolium* hosts, the chromosomes are painted to show only those segments whose parentage is conserved across all isolates from: the PoT lineage, wheat plants, the PoL lineage, *Lolium* plants. Segments are colored according to their respective donors.

Hybrid swarms are characterized as populations in which hybrid individuals undergo subsequent inter-mating and backcrossing to the parents. In some organisms, this can lead to extinction of the parental genotypes and formation of new hybrid species (*34*). The chromopaintings showed clear evidence of genetic recombination within and between hybrid individuals from both the PoT and PoL populations, with the most extreme examples being instances where entire recombinant chromosome structures were shared between PoT and PoL members (e.g. chromosome 1 in PoL2 and PoT3; and chromosome 5 in PoL3 and PoT7). Likewise, many of the crossovers between chromosome segments with different ancestries were widely distributed among the various haplotypes, yet showed abundant evidence of re-assortment and recombination. In fact, a detailed crossover analysis revealed several cases where different haplotypes contained reciprocal crossover products (Fig. S6; Table S3), which can only be reasonably explained if there were abundant inter-mating among sibling progeny arising from single meiotic events.

The variable chromosome ancestry patterns (Figure 3) clearly explain the extreme sequence diversity in the PoT/PoL lineages (Figs. 1 & 2), with the most striking examples consisting of entire chromosomes that were inherited from different donors (e.g. chromosome 4 in PoT1 vs. PoT15; and chromosome 7 in PoT1 vs. PoT15). The figure also illustrates very nicely why different strain comparisons yielded variable patterns of sequence divergence along the chromosomes (Fig. 2A, bottom panel), and this translated to considerable variation in the relative proportions of chromosome ancestry among haplotypes (Figure 4A, Table S4). This was especially true for the PoT lineage, where only about 20% of the genome had the same heritage across all members, and this fraction was even lower for those haplotypes that were found on wheat (Figure 3B).

**Figure 4.**
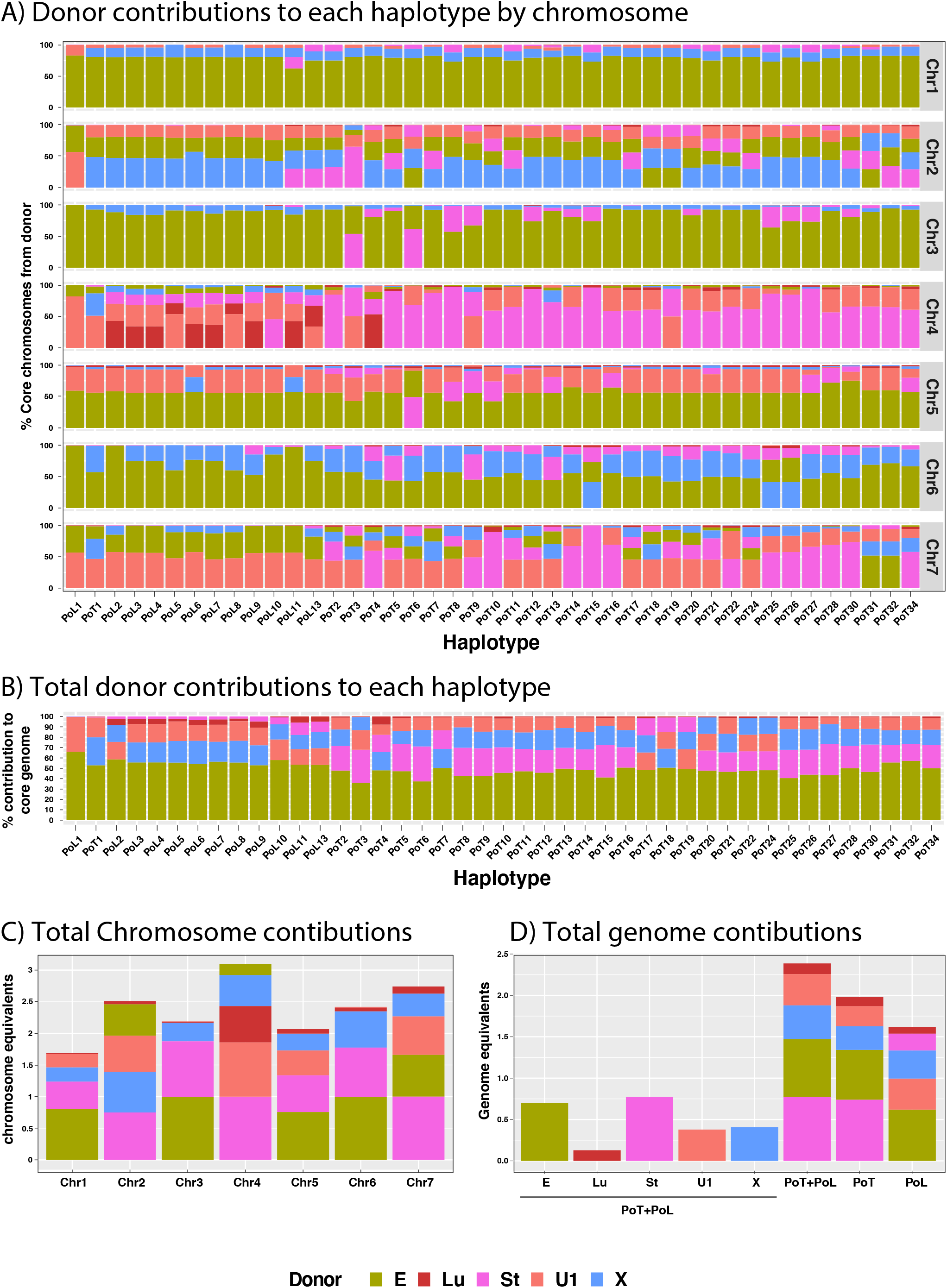
Proportions of PoT/PoL genomes contributed by the respective donor lineages. Chromosome (A) and genome-level (B) contributions to each haplotype were estimated from the chunklengths.out files produced by ChromoPainter. C) Total donor lineage contributions to PoT/PoL by chromosome. The core chromosome contributions from each of the original donor lineages was estimated from the ChromoPainter copyprobs files after filtering out sites whose inferred ancestry was not at least two-times more likely than the next best donor(s). D) shows the total genome equivalents contributed by each donor lineage, and the stacked plots on the right give estimates of pangenome size (in genome equivalents). In all plots, stacking order is from the greatest contribution to least. Only contributions from the main donors are shown (other contributions were less than 1%).

### Swarm reconstruction points to recombination through sexual and parasexual pathways and identifies secondary admixture contributions

Mating in *P. oryzae* typically involves hyphal fusion between two strains of opposite mating type. Therefore, to explain the chromosomal constitutions of PoT/PoL, we hypothesized that their evolution must have occurred in a step-wise fashion involving sequential matings. Specifically, we posited that the process started when two fungal individuals from separate populations with histories of divergent evolution underwent a mating which resulted in the establishment of an initial recombinant haplotype. A member of this founding haplotype then mated with a member of a third diverged population, and so on. At each stage of the process, inter-mating among members of the burgeoning population created a hybrid swarm whose collective activities created the present-day PoT/PoL population(s). Furthermore, because all chromosome segments with shared ancestry showed extremely low sequence divergence (Figures 2A, C & D), we predicted that the founder event must have happened quite recently - maybe not long before blast was first reported on ryegrass in 1971.

i) *The first admixture event that precipitated WB/GLS evolution occurred in approximately 1958.* To estimate the date of the first hybridization event, we took advantage of the fact that all PoT/PoL members co-inherited 1 Mb sequence blocks on chromosomes 1 and 2 from PoE1 and PoU1, respectively (Figs. 3 & S6). These sequences were used for tip-dating to establish a time-frame for the date of initial PoT/PoL divergence from the most recent common ancestor. Using a carefully curated SNP dataset, we arrived at an estimate of 60 years from the date of the most recent strain collection to the most common recent ancestor (1940 - 1973, 95% CI) (Figure 5A), with an average substitution rate of 7.43E-8 SNPs/site/yr. Consistent with a recent evolutionary origin, only 169 SNPs were identified across the two regions (n = 77 isolates), with an average pairwise divergence between isolates of 11 SNPs (max. = 27). A haplotype network built using the tip-dating dataset showed that most SNPs occurred on terminal branches with very few on edges connecting reticulated nodes (Figure 5B). This pattern indicates that virtually all recombination occurred very early on during PoT/PoL diversification but then ceased, with most of the nucleotide divergence occurring over an extended period of strictly clonal propagation. In fact, only one haplotype appeared to be have arisen through recombination in the epidemic population (bold, red edges) because the other reticulations were due to recombination of standing variation (bold lines). That recombination was largely restricted to one major episode is supported by the fact that, among 39 PoT isolates collected within seven years after the initial outbreak, 26 (67%) defined novel haplotypes and, since then, only 7 of 36 isolates (24%) were new.

**Figure 5.**
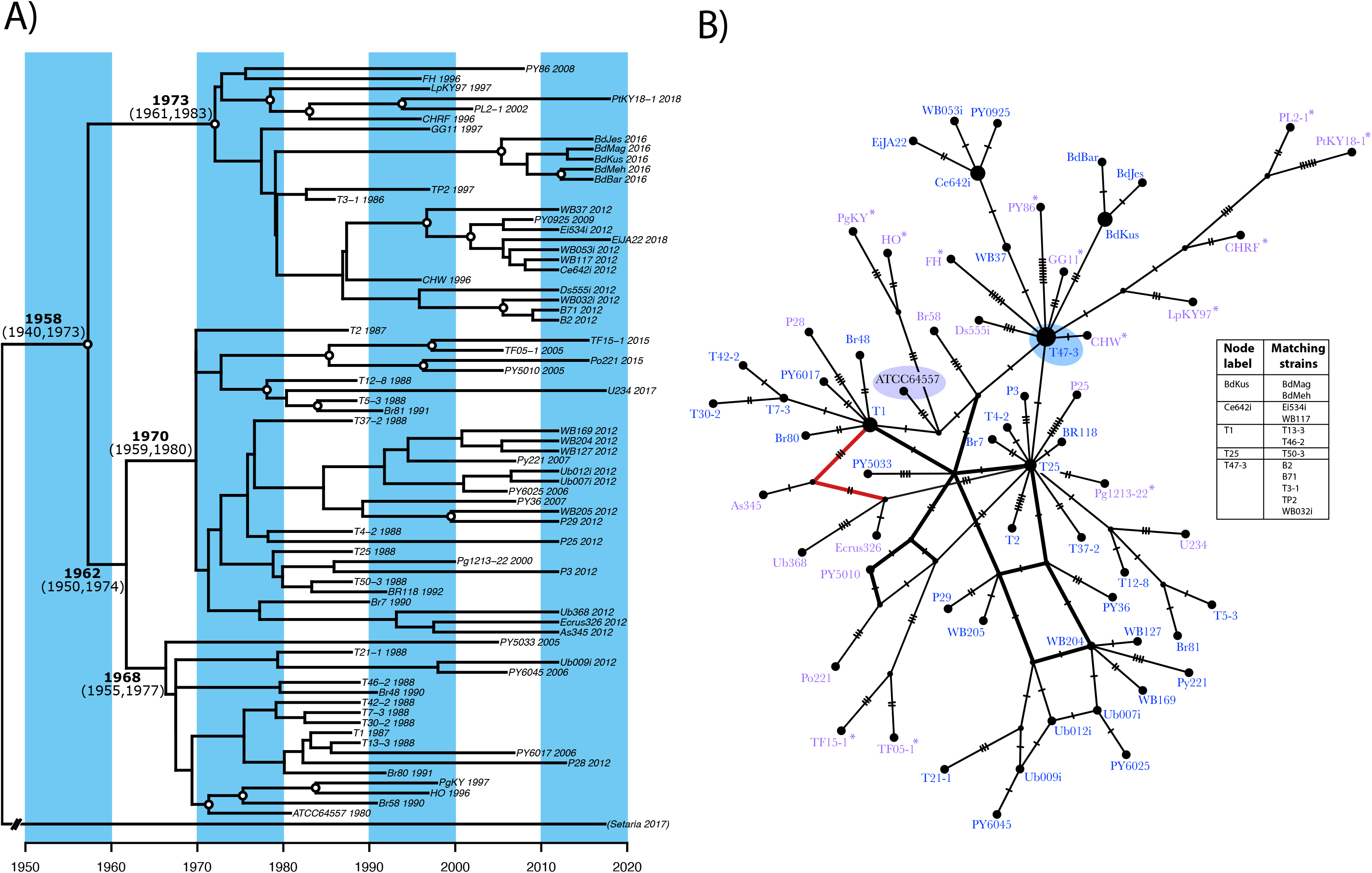
Timing of the first admixture event as inferred by tip-dating using SNP datasets for conserved regions on chromosomes 1 and 2. Estimated dates and 95% confidence intervals are shown adjacent to relevant nodes. White circles on nodes indicate >70% posterior probability. Included is an outgroup isolate from *Setaria* (divergence to ingroup estimated as 18,073 years ago, 95% CI 11,225-25,858). B) Haplotype network for the same SNP dataset using the median spanning network (Bandelt et al. 1999) implemented in PopART (Leigh et al. 2015). The area of each node is proportional to the number of isolates with that haploype, and each tick mark on a connecting edge represents a single SNP. The PoL and PoT founder strains are highlighted with gray and purple ellipses, respectively. Thick black lines are used on edges showing reticulation due to recombination of standing variation (i.e. within the swarm). Thick red lines are used on edges indicating gene flow in the epidemic population (i.e. recombination of new mutations).

ii) A fungal strain isolated from annual ryegrass five years before the first wheat blast outbreak is a descendent of the original hybrid that founded WB/GLS. Nearly all PoL isolates were collected during the mid-1990’s epidemics, with the exception of strain ATCC64557 which was isolated in 1980 and immediately deposited in the American Type Culture Collection. Chromopainting revealed ATCC64557 (PoL haplotype 1; PoL1) to be a two-way admixture (Figure 3) and, based on the proportional inheritance of donor DNAs (∼60:40) (Figure 4B, Table S4), likely descended from an F1 progeny of a mating between strains from the PoE1 and PoU1 populations (Figure 6, admix #1). Critically, with the exception of a few secondary PoE1 introgressions in some haplotypes (see below), the PoE1 and PoU1 segments present in ATCC64557 account for all PoE1 and PoU1 DNA in the PoT/PoL populations (Figure 3). This, along with the absolute conservation of certain PoE1-PoU1 crossover points (fig. S6), led to the confident conclusion that the PoL1 haplotype represents the first step in PoT/PoL evolution and was the original founder to all PoT/PoL strains (Figure 6).

**Figure 6.**
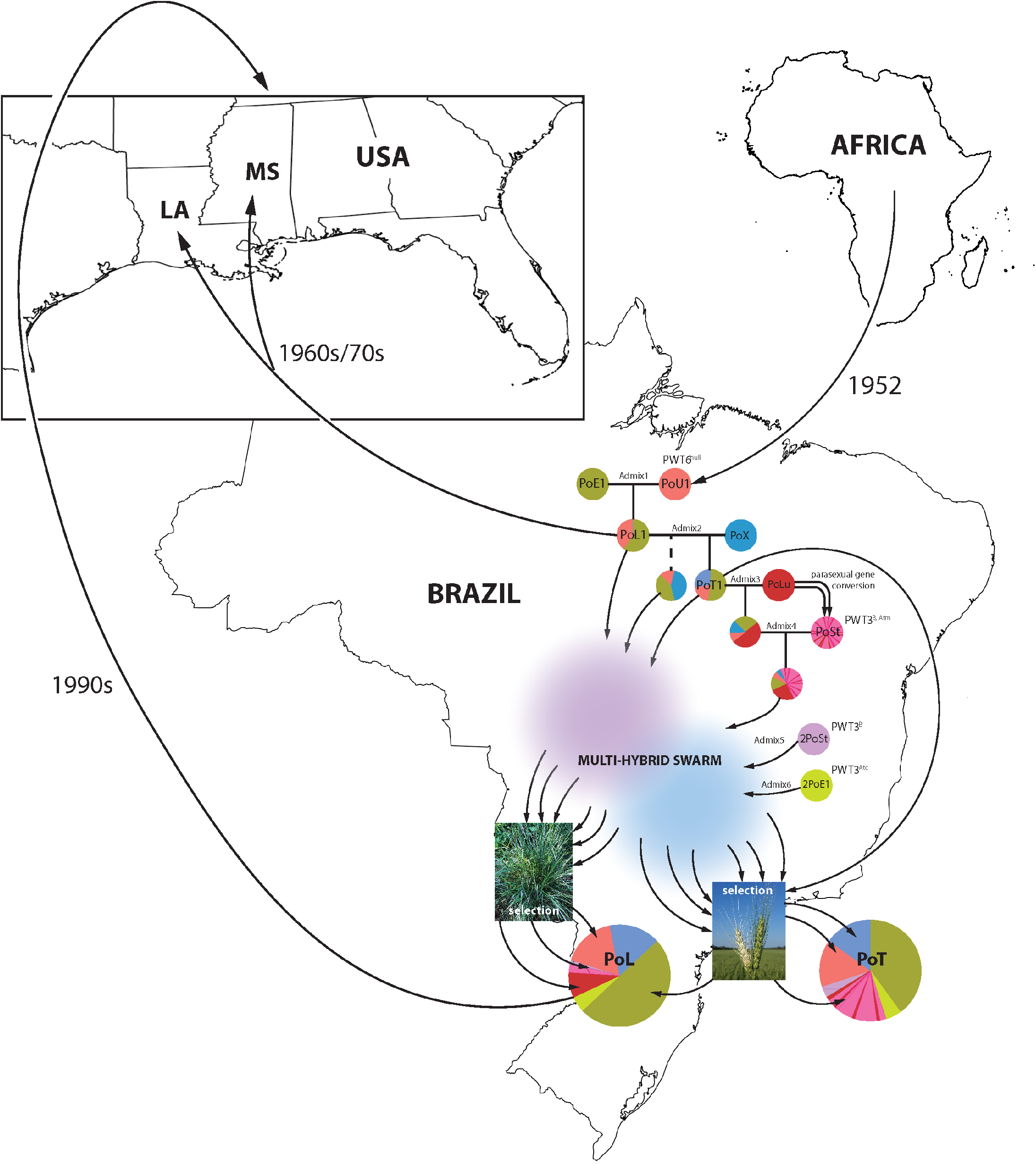
Multi-hybrid swarm model for wheat blast/gray leaf spot evolution. Admixture #1 occurred after PoU1 was introduced into Brazil from Africa in 1952. The PoL1 founder lineage was subsequently introduced into the USA and caused outbreaks in LA and MS. A PoL1 member then initiated a series of sexual/parasexual crosses with isolates specialized on different host grasses that stacked a set of core introgressions through a single filial lineage. This produced a set of “gateway” haplotypes that initiated the swarm activity with the different parental generations and possibly sibling progeny. As spores escaped the swarm they experienced differential selection on wheat and *Lolium* grasses, resulting in lineages that are resolved by host affinity. New haplotypes pathogenic to *Lolium* were then re-introduced into the U.S.

iii) *The first two wheat blast isolates ever recovered are clonal descendants of a hybrid between PoL1 and a member of PoX.* Among the available WB collections, we identified a number of strains collected between 1985 and 1988. Whole genome sequence comparisons and chromopainting revealed that strains T47-3 and T3-1 (from 1985 and 1986, respectively) are essentially clones of one another and constitute a 3-way admixture, with about 27% of their genomes having been contributed by PoX, and the remainder essentially being the PoL1 haplotype. This seemed to confirm a stepwise pattern to WB/GLS evolution, with the second step being a mating between a member of the PoL1 founder haplotype and a PoX isolate (Figure 6, admix #2). Although the resulting haplotype was named PoT1 to reflect the fact that it was the first haplotype to be isolated from wheat, strictly speaking, it groups phylogenetically with PoL (Fig. S1). The PoX introgressions in PoT1 account for all PoX DNA found in PoT members, and ∼65% of that found in PoL. This, along with the widespread conservation of crossover points between PoX segments and segments inherited from PoE and PoU1, provides a strong indication that PoT1 is the founder to all PoT haplotypes and a majority contributor to PoL. The additional PoX DNA seen in some PoL haplotypes (Fig. S6), is most likely from a sibling progeny from admixture #2 (Figure 6).

iv) *PoT and PoL acquired PoLu introgressions via different mechanisms.* At first, the chromosomal haplotypes of PoT and PoL pointed to a divergence in evolutionary pathways following admixture #2, with PoT having acquired PoSt DNA, and PoL gaining the PoLu sequences on chromosome 4. However, closer inspection revealed that all PoT haplotypes contain short blocks of PoLu DNA on several chromosomes and some PoL have PoSt sequences. The PoLu sequences that are found in PoT are especially noticeable on chromosome 4 (fig. S6), where they interrupt long PoSt introgressions. The short lengths of the PoLu tracts, their spacing, and the relatively low background crossover frequency/high introgression length, makes it unlikely that they are crossover products. Instead, they probably arose via gene conversion, which is less prone to interference (*35*). Four of the tracts were >10 kb in length (largest 25.5 kb, Table S5), which is much longer than what is typical for meiotic conversions in fungi (<1 kb) (*36, 37*) and more in line with mitotic conversions (*37–39*). This strongly suggests that the PoLu sequences in PoT were acquired in parasexual crosses that occur after fusion of vegetative hyphae (*40*). Long PoLu conversion tracts were found on the other chromosomes and, for those not inherited from PoT1, all were located in PoSt introgressions, which suggests that the a PoLu isolate underwent a parasexual cross with a PoSt donor, and a subsequent donation to the swarm was made by a progeny isolate (Figure 6). In contrast, PoLu sequences were largely restricted to chromosome 4 in PoL, and the absence of conversion tracts elsewhere suggests that this contribution was acquired in a sexual cross.

At first glance PoSt sequences appeared to be largely absent from the PoL lineage which suggested that it diverged from PoT before the donation of PoSt sequences. The small number of PoSt introgressions potentially could have been acquired via subsequent gene flow. However, nearly all the haplotypes with PoSt sequences (e.g. PoLs 5, 6, 7, 8, 9, 10 and 13) showed no evidence of gene flow in the haplotype network (Figure 5B). In addition, PoT haplotype #4 not only contained admixture contributions from all known donors but it also had the longest PoLu introgression. This was also hard reconcile with gene flow post-divergence and led us to suspect that PoT4 might be a lone holdout from a population of “gateway” haplotypes that originally contained contributions from all donors, and from which PoT and PoL diverged through the differential loss of admixture contributions. To test this possibility, we used reciprocal crossover analysis to identify residual PoSt sequences in PoL that might have belonged to larger introgressions. One example was found on chromosome 3 in PoL6 and PoL7 that appeared to have broken up by a crossover whose reciprocal product segregated into haplotypes PoT4, PoT14, PoT20, PoT30, and PoT31 (fig. S6; Table S3). Not only is this in line with elimination of PoSt sequences from PoL but, the fact that PoT4 was a recipient of the other crossover product, supports the idea that it, and possibly other fully admixed individuals, was a critical nexus in PoT/PoL evolution. Further evidence that PoL divergence involved PoSt sequence elimination comes from the fact that even PoL strains without visible PoSt contributions in Figure 3 have several short (meiotic) conversions peppered throughout their genomes, and these occur in chromosomal positions where some PoT haplotypes have PoSt sequences (data not shown). Most of the original PoSt donations were probably eliminated when PoL members backcrossed with PoL1, PoT1 and/or a meiotic sibling of PoT1 - occurrences that are evident in figure S6.

For the reciprocal elimination of PoLu sequences in PoT, it appears these sequences were evicted when a PoSt admixture, as PoT15 inherited chromosome 4 in its entirety from a PoSt donor (Figure 3). Backcrossing to PoT1 and subsequent inter-mating would have then produced the reduced-PoSt chromosome 4 configurations seen in the other PoT haplotypes.

*The swarm received donations from two different members of the PoSt lineage, with timings that likely predated wheat blast’s initial emergence.* As noted above, the pattern of PoLu conversions in PoT point to admixture with a PoSt member prior to the latter’s donation to the swarm. However, the PoSt introgressions in some PoT haplotypes lacked these conversions, pointing to another donation from a second PoSt isolate. This would also explain why the fraction of the PoSt genome found in PoT/PoL was ∼ 75% (Figure 4D, Table S6), instead of the 50% expected for a single cross. We confirmed this suspicion by examining regions of chromosomes 4 and 7 that showed PoSt heritage, which identified two distinct PoSt haplotypes (Fig. S7A & C; and data not shown). One haplotype was identical to the *Urochloa* pathogen, Up35 (a PoSt member according to DAPC, Fig. S2), while the other matched *bona fide Stenotaphrum* pathogens, exemplified by strain U233. The Up35-like PoSt contribution was already present in one of two isolates collected in 1986 (T49-1) and all isolates henceforth, which effectively dated the first PoSt donation prior to the 1986 wheat growing season and, by extension, the PoLu introgression on chromosome 4 prior to that.

vi) *Chromosomal pedigree analysis revealed a second contribution from the PoE1 lineage.* Several PoL members contained PoE1 introgressions that are not present in PoT1 and, therefore, appeared to have been acquired by backcrossing with the PoL1 founder (Figure S6). A good example is on the left arm of chromosome 7 where the reacquisition of a diagnostic PoE1 to PoU1 crossover strongly supports the occurrence of a backcross. Several PoT haplotypes also contained PoE1 introgressions that were not present in PoT1. At first, we assumed that these segments were acquired via the same route. However, critical PoE1 segments on chromosomes 5 and 7 could not have come from backcrosses as they were in regions where PoL1 showed PoU1 heritage (Fig. S6), which instead told us that these segments must have originated in a second PoE1 donor (2PoE1). This was subsequently confirmed through a detailed SNP analysis which identified several chromosome segments with haplotypes that were very different to PoL1 but showed high identity to other PoE1 members, with the closest similarity being to strains U169 and U229 from Uruguay. (fig. S7B & D). 2PoE1 sequences were not detected in PoL members (data not shown) which could mean that they never received these donations but, more likely, such contributions - along with the PoSt sequences - were purged by backcrossing with PoL1.

vii) *The accumulation of Repeat Induced Point mutations (RIP) between admixture generations confirms a role for the sexual cycle in driving PoT/PoL diversification.* It seemed likely that the abundant recombination was at least partially driven by the sexual cycle, which, although easily established in the laboratory (*41–43*), has never been observed in nature. To test for sexual reproduction, we scanned the PoT/PoL genomes for evidence of recent <u>R</u>epeat Induced <u>P</u>oint mutation (RIP) events. RIP is a sexual cycle-specific process in fungi that was first discovered in *Neurospora crassa* (*44*) and then in other fungi, including *Pyricularia* (*45*). It is activated during the sexual cycle and causes mutational sweeps comprising many C to T transitions on both strands of a DNA duplex either preceding, or during, the last round of pre-meiotic DNA synthesis (*46*) (Figure 7A). To test for a role for the sexual cycle in PoT/PoL evolution we looked for long, asymmetric RIP tracts that occurred subsequent to PoT1’s origin. Whole genome alignments of T47-3 against T1-1 and T2-1 (collected in 1987) identified several exceptionally long tracts of *de novo*, unidirectional RIP mutations (Figure 7B), which suggests that these isolates are possibly F1 sexual progeny of a PoT1 isolate. Raw haplotype data were used to confirm the T47-3 heritage of these sequences to rule out inheritance of pre-RIP’d sequences from an admix donor.

**Figure 7.**
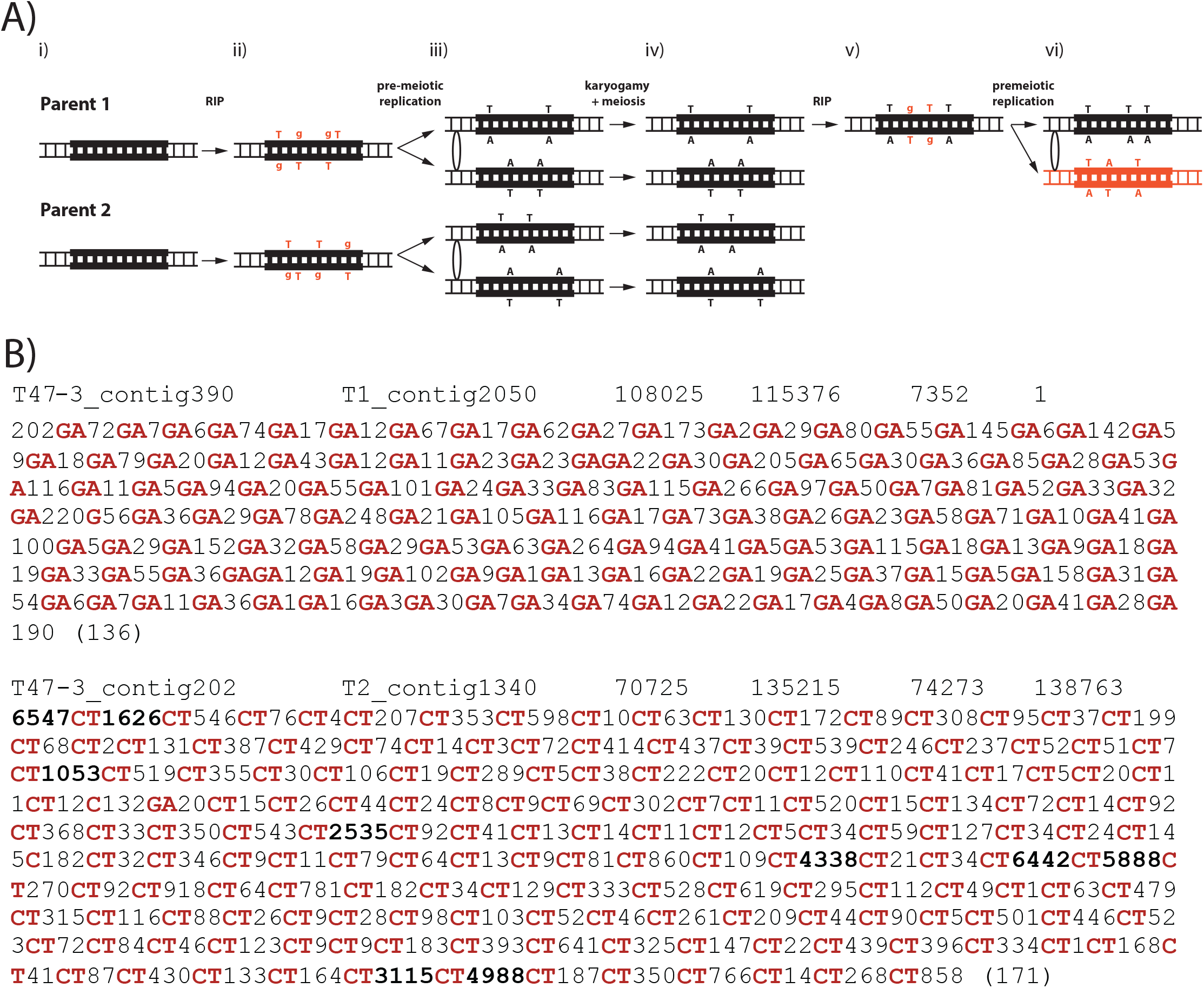
*De novo* RIP mutations provide evidence of sexual activity in the swarm. A) RIP causes unidirectional C to T transitions in a single sexual cycle. i) DNA strands containing a repeated sequence (bold lines) in two parental isolates experience independent RIP events affecting both strands of the DNA duplex just before, or during, pre-meiotic DNA replication. A.ii) T nucleotides resulting from RIP become mismatched with Gs on the opposite DNA strands; A.iii) premeiotic DNA replication resolves the mismatched bases, and the mutations then segregate into the 4 meiotic products (A.iv) producing uni-directional RIP patterns; A.v) RIP in a subsequent cross will produce a bi-directional RIP pattern in the daughter molecule (shown in red). B) Examples of *de novo* RIP mutations in wheat blast isolates T1-1 and T2-1 (both from 1987). Shown are blast alignments (-evalue 1e-50, -outfmt 6, using the btop - traceback operations option). Columns are as follows: qseqid, sseqid, qstart, qend, sstart, send, followed by the btop string, which shows # of aligned bases, nucleotide in query, nucleotide in subject, # of aligned bases, etc. RIP mutations are shown in red, and long (> 1 kb) blocks of unmutated sequence between RIP’d regions are highlighted in bold.

viii) *PoT/PoL divergence was driven by structured mating within the swarm.* The clear phylogenetic separation of PoT from PoL, as well as their distinct chromosome configurations would lead to the logical conclusion that their divergence came about as a result of host adaptation. However, the lack of reticulation in the haplotype network (Figure 5B) implies that once host infection occurred, mating activity ceased, presumably because access to the swarm was lost. More importantly, the trends in chromosome configurations that distinguished PoT from PoL covered so much of the genome, and were so variable within the lineages, that it is hard to formulate a host-driven selection scenario that would account for the differences. As just one example, we would have to infer that *Lolium* imposes a selection against virtually all of the PoSt sequences on all chromosomes (see Figure 3). Instead, we conclude that the divergence in chromosome configurations is largely coincidental and mostly due to structure within the swarm that caused the two lineages to have differential access to potential mating partners.

### Sequential admixture drove broad infection capability on wheat through multiple, independent contributions of null/non-functional alleles for two different host-specificity genes

Prior work has shown that successful infection of wheat requires the loss or inactivation of the *PWT6* and *PWT3* host-specificity genes that would normally trigger resistance in wheat containing the corresponding resistance genes (*29, 47*). Therefore, we sought to understand the dynamics of these genes in the context of PoT/PoL evolution.

i) *PWT6 was deleted in the PoU1 lineage and was donated to PoT/PoL in admixture 1*. *PWT6* is present in *P. oryzae* isolates from *Eleusine* and *Eragrostis*, yet absent from PoL, which prompted the conclusion that the gene was lost when PoL branched from the *Eleusine* lineage (*47*). However, because our data show that PoL evolved through lineage convergence and not branching, we suspected that the *PWT6* was already missing in the PoU1 lineage and donated to PoL in admixture 1. Using chromosome-level, reference-guided genome assemblies, we first determined that the correct chromosomal position for PWT6, and the corresponding null locus, is in the middle of chromosome 5 (fig. S8A), and not the end of chromosome 3 as originally reported (*47*). A quick inspection of PoL1’s haplotype at this locus told us that the donor was probably the PoU1 parent and, consistent with this prediction, all PoU1 isolates were found to lack *PWT6*. A detailed analysis of sequence composition and chromosome organization spanning the deletion breakpoints in these isolates revealed perfect nucleotide and structural identity to the PoL1 *pwt6^null^* locus (fig. S8A), with a copy of the repetitive 5S RNA gene defining the breakpoints (fig. S8B). The chromopaintings revealed that this allele was widely present in PoT/PoL but a few PoT haplotypes were predicted to have acquired the relevant chromosome segment from the PoSt donor (Figure 3 & S6). This was confirmed by analyzing genome assemblies for relevant PoT isolates and the PoSt members, Up35 and UbJA112, which revealed that they have the same rearrangement with a pair of indels flanking the breakpoints (fig. S8A).

ii) *pwt3^B^ was contributed to the swarm in the PoSt admixture.* Inoue and coworkers previously recognized that one of the non-functional alleles in PoT, *pwt3^B^*, was on a chromosome segment with strong sequence similarity to the *Urochloa* pathogen, Br35 (renamed Up35 in the present study), and they concluded that the allele was inherited from a related isolate (*29*). This conclusion is supported by our finding that all isolates possessing *pwt3^B^* inherited the relevant chromosome segment from the PoSt lineage (asterisks, fig. S6), of which Up35 is a member. *pwt3^B^* had already reached an appreciable frequency just three years after the initial outbreak, being present in 3 of 7 isolates analyzed from 1988 (Table S7). However, because PoSt introgressions occurred prior to 1986 (see above), we surmise that *pwt3^B^* was already present in the initial “outbreak” population.

iii) *pwt3^Atc^ was contributed to the swarm by the 2PoE1 donor.* The *pwt3^Atc^* allele reported by Inoue and coworkers had a compound insertion of three different transposons in the coding sequence (*29*). Using BLAST searches to survey the non-PoT/PoL isolates in our study, we found identical transposon insertions in five isolates from *Eleusine*, including the isolate most closely related to the 2PoE1 donor. Chromosome ancestry and detailed haplotype analyses confirmed that *pwt3^Atc^* resided on a 2PoE1 introgression segment (fig. S6). *pwt3^Atc^* now appears to be the predominant allele in the present-day wheat blast population (Table S7) but was first detected in 1988 in haplotype PoT4.

iv) *pwt3^Atm^ arose via a de novo transposon insertion that occurred after the PoSt admixture event.* The *pwt3^Atm^* allele contains an MGL retrotransposon insertion in the promoter region and was already present in the two haplotypes collected in 1987 (PoT2 and PoT3) (Table S7). Flanking sequence analysis revealed that the insertion occurred in a PoU1 allele and, because all PoT isolates descended from PoT1, which possesses the PoU1-derived *PWT3^A^*, this told us that *pwt3^Atm^* must have arisen through a *de novo* transposon insertion that occurred after admixture 2. Initially, this finding seemed to support the springboard hypothesis, which posits that blast initially became established on *rwt3* wheat which increased the likelihood of MGL insertion in *PWT3*. However, due to the lack of explosive propagation that should have come about due to the sudden niche expansion, we suspected that the MGL insertion might have happened after the PoSt (and *pwt3^B^*) introgression had already occurred. To explore this possibility, we leveraged the fact that most repeated sequences experience independent RIP mutations in each mating, which opened up the possibility of exploring whether the MGL copy in *PWT3* originated in the PoT1 genome or was introgressed from the PoSt donor (figs. S8C & D). Interrogation of Illumina assemblies for all isolates containing *pwt3^Atm^* identified three strains (T12-8, T46-2 and Br81) where the assembly extended from the *pwt3* sequences well into the MGL element. This revealed that the *pwt3^Atm^* insertion was identical to a copy found in the *Urochloa* pathogen, Up35, apart from an abundance of G to A and C to T transition mutations that affected the element inserted in *pwt3^Atm^* (fig. S8E). There were some rare RIP-type mutations in the most closely related element in Up35 but this may be because this strain possesses a small number of mutations relative to the actual PoSt donor. In contrast, the MGL in *pwt3^Atm^* was highly divergent relative to the closest sequence(s) in the PoT1 isolate T3-1, with numerous RIP mutations being present in the T3-1 copy (fig. S8F). This effectively ruled out the possibility that the insertion happened prior to admixture 3 because the null hypothesis holds that T3-1 should not have undergone another sexual cycle. To ensure that a more related element was not overlooked due to incomplete assembly of repeated sequences, we confirmed the absence of diagnostic sequence motifs by interrogating raw sequence reads for the diagnostic mutations.

v) *The pwt3^Atp^ allele arose through a recent transposon insertion and is associated with a host jump from Lolium onto U.S. wheat after wheat blast was fully established in Brazil.* The *pwt3^Atp^* allele contains a Pot3 (MGR586) transposon insertion in the promoter and was found only in isolate WBKY11 that was collected from a diseased wheat plant in the USA in 2011 (Table S7) (*48*). WBKY11 belongs to PoL5 - a haplotype whose other members are all *Lolium* pathogens and possess *PWT3^A^.* Given that these isolates are related to WBKY11 by clonal descent, this identifies *pwt3^Atp^* as a second *de novo* insertion event. It seems likely that this mutation was a critical factor in allowing WBKY11 - ostensibly a *Lolium* pathogen - to jump hosts.

## DISCUSSION

There is now compelling evidence that ancestral alleles and gene flow play important roles in the adaptation/speciation of a range or organisms (*8*). Clearly, when rapid responses to new ecological opportunities or perturbations are required, the ability to re-utilize/re-purpose standing variation would be more beneficial that waiting for *de novo* mutations to arise. How novel combinations of old alleles are formed, however, and their genetic architectures within newly-evolved populations is still poorly understood. Likewise, the relative contributions of pre-existing variants versus new mutations in driving adaptation are unknown. Here, we shed new light on these questions with a comprehensive reconstruction of a recent radiation in the blast fungus. Ours is a landmark achievement because we were able to document the entire process, starting with the identification of all sources of standing variation, through the establishment of mechanisms driving diversification, the genetic architecture of the newly-evolved population, and the functional consequences thereof. It could be argued that the events we characterized do not fit into the standard definition of an adaptive radiation because the gene flow was between allopatric populations of the same species, and it did not result in the generation of new species. Here, we would simply point out that the mechanisms that drive evolutionary processes almost certainly don’t draw such distinctions.

Adaptive radiations are thought to occur in response to novel ecological opportunities (*49*) but that does not appear to be the case for wheat blast or gray leaf spot because wheat and *Lolium* have been grown in Brazil since the end of the 19th century (*50*). Likewise, *P. oryzae* is globally distributed in regions with long histories of wheat production and/or *Lolium* presence. Instead, it appears that the process was initiated by sexual opportunity afforded by a chance encounter between fungal individuals from the *Urochloa*- and *Eleusine*-infecting lineages. Curiously, although the founder mating produced a lineage that until now has only been detected in the USA, *P. oryzae* is not known to occur on *Eleusine* or *Urochloa* in N. America, despite ongoing surveillance (M. Farman, personal observations). On the other hand, *Eleusine*-infecting *P. oryzae* lineages are endemic in Brazil and, because molecular dating places the founder cross shortly after 1954, when Brazil first imported African *Urochloa* as a forage to support the beef industry (*51*), it seems plausible that co-introduction of the pathogen then precipitated a cascade of events that now threaten global wheat production. How PoL1 subsequently came to cause disease on *Lolium* in the U.S. is unclear but it may have been introduced from S. America on infected seed, or was possibly swept in when Hurricane Edith arrived from central America just three months prior to the 1971 disease outbreaks.

Rather fittingly, thinking on wheat blast evolution has paralleled that in evolutionary biology. In line with early theory, prior data were interpreted according to a model where host-specialized lineages differentiated in a bifurcating fashion through the gradual and step-wise accumulation of mutations in host-specificity factors (*47*). The differentiation of *Lolium* pathotypes from the PoE lineage was associated with the loss of *PWT6*, and the subsequent evolution of wheat blast was proposed to have involved an initial host jump by a wheat-adapted *Lolium* pathogen, followed by further adaptations involving mutations in *PWT3* (*29*). However, just as modern theory has come to consider standing variation and admixture more prominent players in organismal evolution, we now show that the wheat and *Lolium* pathogens actually co-evolved in a radiation fueled almost entirely by the recombination of standing variation present in five host-specialized lineages. *De novo* mutation seems to have played little, if any, role in the adaptive process because most of the key mutations in host-specificity factors were from standing variation, and few *de novo* mutations arose between the founder event and disease emergence (an estimated 60 SNPs per strain) and most were isolate-specific (Figure 5B). Even the *de novo* transposon insertions in *pwt3^Atm^and pwt3^Atp^* are of questionable significance owing to their redundancy with other *pwt3* alleles segregating in the swarm.

Current thinking in pathogen evolution supposes that host jumps lead to pathogen diversification (*52*). Not only do our data show that the opposite is possible - that diversification can lead to host range expansion - but they also support a model of saltational evolution with new host acquisition being driven purely by chance encounters and random recombination events that produced drastic and highly consequential changes in both phenotype and genotype, over a very small number of generations. First, we found no indication of an initial host jump by a *Lolium* pathogen because the only *Lolium*-infecting haplotype known to exist before wheat blast’s emergence (PoL1) has never been found on wheat, and the other PoL haplotypes’ chromosome configurations clearly identify them as descendants of - and not ancestors to - the wheat blast founder, PoT1, which conversely has not been found on *Lolium*. Thus, the present-day *Lolium* pathogens seem to have re-adapted to the host - possibly expanding their compatibility to include additional species (*L. perenne* and *L. arundinaceum* syn. *Festuca arundinacea*). On the other hand, all isolates from wheat minimally possessed PoX introgressions, suggesting that base-level compatibility to wheat was acquired in admixture #2, possibly due to the introduction of a key virulence factor (or another non-functional avirulence gene). This would explain why a PoX chromosome segment harbors the only diagnostic marker (C17) that reliably distinguishes the PoT and PoL lineages (*53*) despite the extensive recombination (fig. S6).

It is also questionable as to whether *rwt3* wheat served as a springboard for the evolution of *pwt3* mutants as originally proposed (*29*), because there was no indication that any one haplotype experienced a dramatic expansion before the admixtures contributing non-functional *pwt3* alleles appeared and came to dominate the landscape. This, along with the steep trajectories of *pwt3* allele appearances, indicates that these genes were contributed very shortly after PoT1’s genesis and probably before the wheat host was ever encountered (Figure 6). Instead, our data best support a scenario in which admixtures 2 through 6 occurred in quick succession to produce a multi-hybrid swarm that was, quite literally, a “swarm,” with all key donors and their progeny coexisting in a small area, so that a single, fairly brief episode of rampant sexual/parasexual activity resulted in rapid and wide sharing of *pwt3^B^*, *pwt3^Atc^* and *pwt3^Atm^* among PoT members and essentially generated a population that was pre-adapted to wheat infection. What is most remarkable is that, despite the abundant sexual activity within the swarm, almost no recombination was detected after strains started accumulating new mutations (Figure 5B). In other words, mating in the epidemic population appears to be exceedingly rare, and this may be partially related to the low frequency of the *MAT1-2* mating type allele (0.135, this study; 0.1 in ref. (*54*)). This conclusion is supported by the observation that ongoing sampling identifies new PoT haplotypes at an ever-decreasing rate - as would be expected if a one-time set of recombinants were produced in a single, historical event.

Finally, if there is any doubt remaining that wheat blast could have arisen entirely through the spontaneous and random repartitioning of standing variation, with no driving selection from the host, one only has to recall that, in parallel, the swarm generated genotypes capable of infecting at least ten additional grass genera, with some being pathogenic to multiple hosts. Not only does this makes the idea of intermediate host selection seem implausible but it also raises the question as to where crossing occurred such that (non-wheat) host-specific haplotypes (fig. S1) were still recovered? Here we suggest that mating occurred on dead, or dying, host tissues - possibly hay made from a mixture of infected pasture grasses and weeds - as this would have allowed the swarm to generate an untested collection of what might best be described as “hopeful monsters” (though not strictly *sensu* Goldschmidt, (*55*)), which then experienced host-driven selection after their “escape” from the swarm.

When one considers the genetic and pathogenic diversity among the donor isolates, it is not unreasonable to suppose that random mating could lead to the spontaneous generation of a whole array of new host-adapted lineages. In *P. oryzae*, as with other Pyrenomycetes, a single fertilization event can initiate hundreds of independent meioses, as the dikaryon undergoes iterative cycles of replication then karyogamy (see Farman, 2002). This means that, depending on fertility, the last admixture event of the series alone could generate several hundred recombinant genotypes, and with subsequent sib-, inter- and backcrosses, this could easily scale to several million, especially when one considers that sib-mating among germinated ascospores is rapid and common in *P. oryzae* (M. Farman, personal observations).

Theory predicts that when a new population is founded by a small number of individuals, it will have low genetic diversity and, thus, a constrained adaptive capacity (*2*). This would certainly have been true had a *Lolium* pathogen simply jumped hosts. However, because the multi-hybrid swarm incorporated genetic material from five lineages that were themselves highly diverged, this effectively cheated the founder effect and allowed *P. oryzae* to generate a new lineage with greater diversity and, hence, adaptive capacity in ∼20 years, than is present in existing host specialized lineages with histories spanning millenia (and possibly longer) (*30, 31*). With this in mind, it may be no coincidence that so many newly-emerged fungal diseases have their origins in admixture (*56–59*); and while *P. oryzae* is unique in having cheated the founder effect by collating genetic material from multiple con-specific individuals, in other pathosystems diversification was most often accomplished through interspecific hybridization between two individuals.

While the recombination of several diverged genomes will have helped to overcome the founder effect, it raises other potential barriers to evolution in the form of reduced fitness caused by the segregation of host-specificity factors (*60, 61*), or inherent genomic conflicts due to Bateson-Dobzhansky-Müller (BDM) incompatibilities (*1, 62, 63*). It is, therefore, extremely surprising that the the wheat blast and *Lolium* pathogens retained compatibility to their respective hosts when ∼80% of their genomes were segregating - for as many as four alleles. Likewise, the reintroduction of hundreds of thousands of alleles with divergent evolutionary trajectories did not prevent the respective isolates from producing asexual spores, surviving environmental dispersal, completing at least one full life cycle in the host, and then surviving desiccation, long-term frozen storage, and re-cultivation. This is not to say that host incompatibility and BDM effects do not occur and, in fact, a combination of these influences probably explain why individuals representing reciprocal meiotic products were not recovered. It just illustrates that these factors may not be as pervasive as is assumed.

In summary, the evolution of wheat blast and gray leaf spot is a story of almost unimaginable happenstance. The idea that that a fungus that has historically been considered asexual could have participated in a six/seven-way cross - essentially a fungal orgy - to generate a population with an uncharacteristically broad host range is, in itself, incomprehensible. That the heritage of 80% of the genome would have no apparent impact on host compatibility, or be immune to BDM effects is highly unexpected; and, finally, the clear phylogenetic/chromosomal partitioning between the PoT and PoL lineages is inexplicable based on models of host selection or admixture order/patterns, leaving random genetic drift as the only reasonable, albeit unlikely, explanation. We believe that this is telling us that many adaptive radiations and, in turn, speciation occurs precisely when organisms happen upon those rare circumstances that allow the breaking of rules that would normally be holding these processes in check. With this in mind, we suggest that multi-hybridization events that have been completely overlooked in the past, may prove to unlie many of the adaptive radiations reported in the literature, and may have escaped consideration because they cannot be properly detected using phylogenetic trees, D-statistics, or standard implementations of admixture detection software (*21*).

Here, one might argue that we should not be extrapolating from the present data to make inferences about speciation. However, it is important to remember that already the wheat blast fungus has been proposed as a new species (*20*) and, while it is clearly premature to draw this conclusion based on the fungus’s very recent evolution via gene flow, recall also that our data conclusively show that the population has remained reproductively isolated. Thus, it is quite possible that we have just witnessed the birth of a new species - only time will tell. Finally, if we consider the structure of the newfound population: we have a set of haplotypes, each having infection capability on a given host(s), and each comprising a set of genotypes experiencing ongoing diversification. Over time, as fine-tuning of host compatibility drives further divergence of some haplotypes, while others go extinct, the PoT/PoL clades will eventually recapitulate the *P. oryzae* species tree, with isolates specialized to the same host occupying multiple, seemingly-independent clades. Indeed, a multi-hybrid swarm origin for the *P. oryzae* species as a whole would provide a satisfying explanation not only for the tree architecture, but would also account for the polyphyletic origin of many host-specialized forms, as well as the abundant evidence of incomplete lineage sorting and gene flow on most of the main branches (*21*).

## METHODS

### Fungal cultures

All strains were genetically purified via single-spore isolation and then cultured in 10 ml liquid complete medium (6 g casamino acids, 6 g yeast extract, 10 g sucrose) for 7 d with shaking.

### NGS library production and sequencing

DNA was prepared using previously described methods (*64*). Most libraries were prepared using the Nextera kit (Illumina, San Diego, CA) and were constructed according to the manufacturer’s instructions, but using a tagmentation step of 60 min. A small number of libraries were prepared using the Kapa HyperPlus kit (Roche, indianapolis, IN), according to the manufacturer’s protocol. Sequences were acquired using MiSeq (at the University of Kentucky Advanced Genetic Technologies Center, or Bluegrass Community and Technical College; and using HiSeq (150 PE) at Novogene Corp Inc. (Sacramento, CA).

### Sequence assembly

Raw reads were trimmed to remove adaptor sequences and poor quality regions using Trimmomatic (PE mode: SLIDINGWINDOW:20:20 MINLEN:80 ILLUMINACLIP: NexteraPE-PE.fa:2:30:10). Paired-end reads passing these filters were assembled using Newbler 2.9 or Velvet 1.2.10 with a range of k-mer values 59-129.

### SNP Calling

We employed a custom perl script <u>iSNPcaller.pl</u> which performs two-passes of a repeat-masking algorithm originally developed for TruMatch (*65*) to ensure that SNPs are not called in regions affected by RIP. Briefly, repeat-masked genomes are aligned using blast (-evalue 1e-20, -max_target_seqs 20000 -outfmt btop) and SNPs are then called in uniquely aligned regions that pass a second repeat-filtering step. iSNPcaller routinely outperforms BWT/GATK SNP calling pipelines which produces ∼30% false calls in fungal genomes (Stewart and Farman, in preparation). iSNPcaller accuracy was tested by identifying SNPs in fully independent genome assemblies derived from the same raw read datasets. This revealed a false SNP call rate of ∼2 x 10^-5^.

For tip-based dating, SNPs were re-called by aligning raw reads (where available) to the B71 reference genome using bowtie and then identifying SNPs using the Genome Analysis Toolkit with the parameters: HaplotypeCaller, -ploidy 1; --emit-ref-confidence GVCF; GenotypeGVCFs default settings; SelectVariants -select-type SNP. Custom scripts were used to filter the resulting VCF files using the following schema: >10 reads covering the site in question; an alternate:reference allele ratio of ≥ 10:1. SNPs passing these pre-qualifiers were cross-referenced against genome alignment data and retained only if the site was “called” in every genome, was in a non-repeated region, and was also identified using iSNPcaller.

### SNP filtering for tip-based dating

Inspection of the SNP calls in conserved segments of chromosomes 1 and 2 showed that some isolates possessed short blocks of DNA introgressed from another donor. Therefore, to avoid dating estimates being unduly influenced by variants acquired by recombination/gene conversion, we interrogated SNP calls for more than 250 isolates from more than 20 host-specialized populations and filtered out SNPs if the variant allele was present in one or more populations known to be donors to the WB/GLS genomes.

### Phylogenetic analyses

Pairwise distance data from iSNPcaller were used to build a neighbor-joining using MEGA (*66*) and the radial tree output format was selected. Phylogenetic analysis of MPG1 and CH7BAC7 were performed by aligning the sequences with MUSCLE 3.8.31 (*67*) and genarting maximum likelihood trees with RAXML8 (*68*) using the GTRGAMMA model, and 100 bootstrap replications. The cophylogeny plot was created in R using the cophylo function (assoc=NULL, rotate=TRUE) in phytools (*69*).

### Identifying population subdivision via Discriminant Analysis of Principle Components (DAPC)

Haplotype data in STRUCTURE format were imported into the BIC/DAPC module in the Poppr package (*70*). Clusters were identified using the find.clusters function (max.n.clust=30, n.iter = 1e6). DAPC was performed over a K-value range from 10 to 25, after retaining 100 principle components. Populations used for subsequent analyses were defined according to the partitioning at K=16, as this value had the lowest Bayesian Information Criterion (BIC) value. Similar population divisions were identified with STRUCTURE (*71*), although the latter program failed to resolve the PoO and PoS clades which were clearly separated, both by DAPC (fig.S3) and by phylogenetic approaches (*21*).

### Assessing chromosome ancestry using ChromoPainter

SNP data were used to call individual chromosomal haplotypes for each strain, retaining only bi-allelic sites. The resulting file was parsed to generate standard ChromoPainter (*72*) input files – one per core chromosome. Finally, the datasets were filtered to remove all SNP positions where any one strain had a missing datapoint. Chromopainter was run separately for a single representative of each PoT/PoL haplotype, using all non-PoT/PoL strains as potential donors, and an even recombination fraction of 7e-9 across all chromosomes. Run parameters included the haploid (-j) and print-out copyprobsperlocus files (-b) options, with each expectation maximization being performed over 10 iterations, while maximizing over copy proportions (-ip option). Donor probability plots and chromosome haplotype plots were generated from the resulting copyprobs files, using custom R programs. A custom perl script was used to calculate genome contributions from the samples.out files, and a custom R code was used for plotting.

### Assessing chromosome ancestry using ShinyHaplotypes

The ChromoPainter input files were used as inputs to the perl script SlideCompare which compared haplotypes in pairwise fashion in sliding windows of 2000 variant sites and a step size of 400. Haplotype divergence was calculated as #SNPs/ variant sites. The output files were then read by the ShinyHaplotypes.R code for interactive plotting.

### Tip-dating using BEAST

Due to the limited number of variant sites differentiating any pair of isolates, the SNP data were analyzed as a single partition. The HKY + I + G site substitution model was utilized as determined to be optimal by PartitionFinder2 (*73*). A strict clock was employed and its rate was estimated in the MCMC. Coalescent Extended Bayesian Skyline was selected as the tree prior to gain insight into the population dynamics over time. Twenty independent MCMC samplings were performed, with each comprising 10 million pre-burnin iterations, 100 million sampled iterations, and data logging every 50,000 iterations. Tracer was used to assess convergence in the combined logfiles, with each file showing effective sample sizes (ESS) of > 200. TreeAnnotator was used to create a maximum clade credibility tree which was plotted using the ape (*74*) and phyloch (https://rdrr.io/github/fmichonneau/phyloch/man/) R packages.

## ACKNOWLEDGMENTS

We thank Yukio Tosa for sharing unpublished genome sequence data; and the following for helpful comments: Sophien Kamoun and Thorsten Langner (The Sainsbury laboratory), Kevin McCluskey (Kansas State University/Bolt Threads), Pierre Gladieux (Institut National de la Recherche Agronomique), Nik Grunwald (Oregon State University), David Weisrock (University of Kentucky); and Larry Trevathan (Animal and Plant Health Inspection Service). This work was supported by the United States Department of Agriculture, Agriculture and Food Research Initiative grant 2013-68004-20378, multistate project NE1602; Agricultural Research Service project 8044-22000-046-00D; Hatch project KY012037; the National Science Foundation, MCB-1716491; and the University of Kentucky College of Agriculture Food and the Environment.

## DATA AVAILABILITY

Genome sequence data are available under various NCBI BioProjects IDs as listed in Table S1. Datasets used for the analyses described herein are available at https:// figshare.com/account/home#/projects/125476

## CODE AVAILABILITY

Custom code used to generate figures are available at https://figshare.com/account/home#/projects/125476

