## Supplementary Information for "Recombination of standing variation in a multi-hybrid swarm drove adaptive radiation in a fungal pathogen and gave rise to two pandemic plant diseases"

Table S1. Fungal isolates used in this study

| **Isolate** | **a.k.a** | **Host of Isolation** | **Country** | **Region** | **Year** | **Ref.** | **Phylogenet. lineage** | **DAPC lineage** | **NCBI accession** |
|---|---|---|---|---|---|---|---|---|---|
| As073 | 12.0.073 | *Avena sativa* (oat) | Brazil | MS | 2012 | (*1*) | Lolium1 | n.d.[A] | SAMN18576974 |
| As321 | 12.0.321 | *A. sativa* | Brazil | MS | 2012 | (*1*) | Lolium1 | n.d. | SAMN18576976 |
| As345 | 12.0.345 | A. sativa | Brazil | MS | 2012 | (*1*) | Lolium1 | n.d. | SAMN07829574 |
| As347 | 12.0.347 | A. sativa | Brazil | MS | 2012 | (*1*) | Lolium1 | n.d. | SAMN18576977 |
| Br58 | | | Brazil | PR | 1990 | (*2*) | Lolium1 | n.d. | SAMD00051172 |
| P28 | P-0028 | *Bromus tectorum (cheat grass)* | Paraguay | IT | 2014 | (*3*) | Lolium1 | n.d. | SAMN05864041 |
| P29 | P-0029 | *B. tectorum* | Paraguay | IT | 2014 | (*3*) | Triticum1 | n.d. | SAMN05898532 |
| Ce642i | 12.0.642i | *Cenchrus echinatus* (buffel grass) | Brazil | PR | 2012 | (*1*) | Triticum1 | n.d. | SAMN07829578 |
| Ce535i | 12.0.535i | *C. echinatus* | Brazil | PR | 2012 | (*1*) | Triticum1 | n.d. | SAMN18576978 |
| MG07 | | *C. ciliaris* | India | Bangalore, KA | - | (*4*) | Urochloa2 | PoU1 | SAMN04217096 |
| Cd88215 | | *Cynodon dactylon* (Bermuda grass) | Philippines | Cabanatuan, NE | 1988 | (*5*) | Cynodon1 | PoC1 | SAMN14167123 |
| Cd88217 | | *C. dactylon* | Philippines | Cabanatuan, NE | 1988 | (*5*) | Cynodon1 | PoC1 | SAMN14167123 |

| | | | | | | | | | |
|---|---|---|---|---|---|---|---|---|---|
| CpJA159 | | *Cynodon plectostachyus* (Bermudagrass) | Brazil | MG | 2018 | This study | Cynodon2 | PoC2 | SAMN19488802 |
| Ds555i | 12.0.555i | *Digitaria sanguinalis* (hairy crabgrass) | Brazil | PR | 2012 | (*1*) | Lolium1 | n.d. | SAMN07829577 |
| Ec88443 | | *Echinochloa colona* (jungle rice) | Philippines | Los Baños, LB | 1988 | (*5*) | Echinochloa | PoEc | SAMN19488808 |
| G22 | | *Eleusine coracana* (finger millet) | Japan | - | 1976 | (*6*) | Eleusine2 | PoE2 | SAMN08009554 |
| JP29 | | *E. coracana* | Japan | - | 1991 | (*7*) | Eleusine2 | PoE2 | SAMN19488809 |
| PH42 | | *E. coracana* | Philippines | - | 1983 | (*6*) | Eleusine2 | PoE2 | SAMN08009570 |
| Z2-1 | | *E. coracana* | Japan | Kagawa | 1977 | (*2*) | Eleusine2 | PoE2 | SAMD00051173 |
| MG03 | | *E. coracana* | India | Bangalore, KA | 2013 | (*4*) | Eleusine2 | PoE2 | SAMN04216994 |
| MG04 | | *E. coracana* | India | Bangalore, KA | 2012 | (*4*) | Eleusine2 | PoE2 | SAMN04216996 |
| MG12 | | *E. coracana* | India | Bangalore, KA | 2013 | (*4*) | Eleusine2 | PoE2 | SAMN04217237 |
| B51 | | *Eleusine indica* (goosegrass) | Bolivia | Quirusillas, SC | 2012 | (*6*) | Eleusine1 | PoE1 | SAMN08009542 |
| Br62 | | *E. indica* | Brazil | - | - | (*8*) | Eleusine1 | PoE1 | SAMEA4029901 |

| | | | | | | | | | |
|---|---|---|---|---|---|---|---|---|---|
| CD156 | | *E. indica* | Ivory Coast | Ferkessedougou, SV | 1989 | (*9*) | Eleusine1 | PoE1 | SAMEA4708261 |
| Ei534i | 12.0.534i | *E. indica* | E. indica | PR | 2012 | (*1*) | Triticum1 | n.d. | SAMN07829576 |
| Ei8303 | EiA8303 | *E. indica* | Philippines | Los Baños, LB | 1984 | (*5*) | Eleusine1 | PoE1 | SAMN19488810 |
| Ei88365 | | *E. indica* | Philippines | Santo Tomas, BTG | 1988 | (*5*) | Eleusine1 | PoE1 | SAMN19488811 |
| Ei8927 | | *E. indica* | Philippines | BUDA | 1989 | (*5*) | Eleusine1 | PoE1 | SAMN19488812 |
| Ei9064 | | *E. indica* | China | FJ | 1996 | (*5*) | Eleusine1 | PoE1 | SAMN19488813 |
| Ei9411 | | *E. indica* | China | FJ | 1990 | (*5*) | Eleusine2 | PoE2 | |
| EiJA22 | | *E. indica* | Brazil | Patos de Minas, MG | 2018 | This study | Triticum1 | n.d. | SAMN19488814 |
| EiJA178 | | *E. indica* | Brazil | Viçosa, MG | 2018 | This study | Cynodon1 | PoC1 | SAMN19488815 |
| EiJA56 | | *E. indica* | Brazil | Viçosa, MG | 2018 | This study | Cynodon1 | PoC1 | SAMN19488816 |
| MZ5-1-6 | | *E. indica* | Japan | - | - | (*10*) | Eleusine2 | PoE2 | SAMD00069327 |
| U229 | | *E. indica* | Uruguay | Valle Alto, TT | 2017 | (*11*) | Eleusine1 | PoE1 | SAMN19488817 |
| U231 | | *E. indica* | Uruguay | Valle Alto, TT | 2017 | (*11*) | Eleusine1 | PoE1 | SAMN19488818 |
| U169 | U169 | *Eleusine spp.* | Uruguay | Río Branco, CL | 2010 | (*11*) | Eleusine1 | PoE1 | SAMN19488819 |
| Ecan194 | 12.0.194 | *Elionurus candidus* | Brazil | MS | 2012 | (*1*) | Triticum | n.d. | SAMN18576975 |
| AR4 | | *Eragrostis curvula* | Japan | - | - | (*12*) | Eleusine2 | PoE2 | SAMN19488820 |

| | | | | | | | | | |
|---|---|---|---|---|---|---|---|---|---|
| | | (weeping lovegrass) | | | | | | | |
| G17 | | *E. curvula* | Japan | - | 1976 | (*9*) | Eleusine1 | PoE1 | SAMN08009553 |
| EtKY19-1 | | *Eragrostis tef* (teff) | USA | KY | 2019 | (*13*) | Eragrostis | PoEr | SAMN13964779 |
| pg1213-2 | | *Lolium arundinaceum* (tall fescue) | USA | GA | 1999/2000 | (*11*) | Lolium1 | PoL | SAMN14603777 |
| pg1213-22 | | *L. arundinacea* | USA | GA | 1999/2000 | (*6*) | Lolium1 | n.d. | SAMN08009569 |
| TF05-1 | | *L. arundinacea* | USA | Lexington, KY | 2005 | (*6*) | Lolium1 | n.d. | SAMN08009576 |
| TF15-1 | | *L. arundinacea* | USA | Lexington, KY | 2015 | This study | Lolium1 | n.d. | SAMN14144147 |
| FPH-2015-44 | | *Hakonechloa macra* (Japanese forest grass) | USA | OH | 2015 | (*11*) | Luziola | PoLu | SAMN14144145 |
| BTBa-B1 | | *Hordeum vulgare* (barley) | Bangladesh | Gazipur | 2016 | (*14*) | Triticum1 | n.d. | SAMEA104190806 |
| BTBa-B2 | | *H. vulgare* | Bangladesh | Gazipur | 2016 | (*14*) | Triticum1 | n.d. | SAMEA104190807 |
| TH0012-rn | TH0012, TH12 | *H. vulgare* | Thailand | - | - | (*9*) | Oryza | PoO | SAMEA3231789 |

| | | | | | | | | | |
|---|---|---|---|---|---|---|---|---|---|
| TH0016 | TH16 | *H. vulgare* | Thailand | - | - | (*9*) | Oryza | PoO | SAMEA3232033 |
| Lh8401 | LhA8401 | *Leersia hexandra* (southern cutgrass) | Philippines | Los Baños, LB | 1984 | (*5*) | Leersia | PoLe | SAMN19488823 |
| Lh88405-2 | Lh88405 | *L. hexandra* | Philippines | Los Baños, LB | 1988 | (*5*) | Leersia | PoLe | SAMN14167125 |
| Lh8844 | | *L. hexandra* | Philippines | Cabanatuan, NE | 1988 | (*5*) | Leersia | PoLe | SAMN19488824 |
| Lc8401 | LcA8401 | *Leptochloa chinensis* (red sprangletop) | Philippines | Los Baños, LB | 1984 | (*5*) | Echinochloa | PoEc | SAMN14144146 |
| ATCC64557 | U49 | *Lolium multiflorum* (annual ryegrass*)* | USA | MS | 1972 | (*15*) | Lolium1 | n.d. | SAMN19488825 |
| PL2-1 | | *L. multiflorum* | USA | Pulaski Co., KY | 2002 | (*6*) | Lolium1 | n.d. | SAMN08009571 |
| PL3-1 | | *L. multiflorum* | USA | Pulaski Co., KY | 2002 | (*6*) | Lolium1 | n.d. | SAMN08009572 |
| Po221 | | *L. multiflorum* | Uruguay | CL | 2015 | This study | Lolium1 | n.d. | SAMN14153273 |
| U234 | | *L. multiflorum* | Uruguay | 18 de Julio, RO | 2017 | (*11*) | Lolium1 | n.d. | SAMN19488826 |
| U237 | | *L. multiflorum* | Uruguay | UEPL, TT | 2017 | (*11*) | Echinochloa | PoEc | SAMN19488828 |
| CHRF | | *Lolium perenne* (perennial ryegrass) | USA | Siler Springs, MD | 1996 | (*6*) | Lolium1 | n.d. | SAMN08009548 |

| | | | | | | | | | |
|---|---|---|---|---|---|---|---|---|---|
| CHW | | *L. perenne* | USA | Severna Park, MD | 1996 | (*6*) | Lolium1 | n.d. | SAMN08009549 |
| FH | | *L. perenne* | USA | Hagerstown, MD | 1997 | (*6*) | Lolium1 | n.d. | SAMN08009551 |
| GG11 | | *L. perenne* | USA | Lexington, KY | 1997 | (*6*) | Lolium1 | n.d. | SAMN08009555 |
| HO | | *L. perenne* | USA | Richmond, PA | 1996 | (*6*) | Lolium1 | n.d. | SAMN08009558 |
| LpKY97 | LpKY97-1 | *L. perenne* | USA | Lexington, KY | 1997 | (*6*) | Lolium1 | n.d. | SAMN08009564 |
| PgKY | PgKY4OV2.1 | *L. perenne* | USA | Lexington, KY | 2000 | (*9*) | Lolium1 | n.d. | SAMEA4029903 |
| PgPA | PgPA18C-02, PGPA | *L. perenne* | USA | PA | 1998 | (*9*) | Lolium1 | n.d. | SAMEA4029904 |
| TP2 | | *L. perenne* | Japan | Tochigi | 1997 | (*10*) | Lolium1 | n.d. | SAMN14151737 |
| Wk3-1 | | *L. perenne* | Japan | Yamaguchi | 1996 | (*16*) | Lolium1 | PoL | SAMN14603776 |
| U168 | | *Luziola peruvianum* (watergrass) | Uruguay | Río Branco, CL | 2010 | (*11*) | Luziola | PoLu | SAMN14603775 |
| U171 | | *L. peruvianum* | Uruguay | Zapata, TT | 2010 | (*11*) | Luziola | PoLu | SAMN19488829 |
| Mr051i | 12.0.051i | *Melinis repens* | Brazil | PR | 2012 | (*1*) | Triticum | n.d. | SAMN18576973 |
| 87-120 | | *Oryza sativa* (rice) | - | - | - | (*6*) | Oryza | PoO | SAMN08377452 |
| FR13 | | *O. sativa* | France | - | 1990 | (*17*) | Oryza | PoO | SAMEA4708258 |
| Guy11 | | *O. sativa* | French Guyana | - | 1988 | (*6*) | Oryza | PoO | SAMN06050151 |
| IA1 | ARB114 | *O. sativa* | USA | AR | 2009 | (*3*) | Oryza | PoO | SAMN08009559 |
| IB49 | ZN61 | *O. sativa* | USA | AR | 1992 | (*3*) | Oryza | PoO | SAMN08009561 |

| | | | | | | | | | |
|---|---|---|---|---|---|---|---|---|---|
| IC17 | ZN57 | *O. sativa* | USA | AR | 1992 | (*3*) | Oryza | PoO | SAMN08009562 |
| IE1K | TM2 | *O. sativa* | USA | AR | 2003 | (*3*) | Oryza | PoO | SAMN08009563 |
| INA168 | | *O. sativa* | Japan | Aichi | 1958 | (*10*) | Oryza | PoO | SAMD00051169 |
| Ken53-33 | | *O. sativa* | Japan | Aichi | 1953 | (*2*) | Oryza | PoO | SAMD00051177 |
| MBSD02 | RMg-Dl | *O. sativa* | India | BR | 2016 | (*18*) | Oryza | PoO | SAMN05425585 |
| ML33 | | *O. sativa* | Mali | - | 1995 | (*6*) | Oryza | PoO | SAMN08009565 |
| P-2 | | *O. sativa* | Japan | Aichi | 1948 | (*2*) | Oryza | PoO | SAMD00051176 |
| P131 | | *O. sativa* | Japan | - | - | (*19*) | Oryza | PoO | SAMN02981399 |
| PH0014-rn | | *O. sativa* | Philippines | - | - | (*9*) | Oryza | PoO | SAMN19488830 |
| SSID116 | | *O. sativa* | USA | - | 1997 | (*11*) | Oryza | PoO | SAMN19488832 |
| TH3 | | *O. sativa* | Thailand | - | ND | (*2*) | Oryza | PoO | SAMD00051175 |
| U75 | | *O. sativa* | Uruguay | TT | 2005 | (*11*) | Oryza | PoO | SAMN19488833 |
| U107 | | *O. sativa* | Uruguay | Charqueada, TT | 2009 | This study | Oryza | PoO | SAMN19488834 |
| Y34 | | *O. sativa* | China | YN | 1982 | (*20*) | Oryza | PoO | SAMN02981398 |
| BTTrp-5 | | *Panicum repens* (torpedograss) | Bangladesh | Gazipur | 2016 | (*14*) | Panicum | PoP | SAMEA104190823 |
| BTTrp-6 | | *P. repens* | Bangladesh | Gazipur | 2017 | (*14*) | Panicum | PoP | SAMEA104190824 |
| Pr8202 | PrA8202 | *P. repens* | Philippines | Los Baños, LB | 1982 | (*5*) | Panicum | PoP | SAMN14603774 |
| Pr88165 | | *P. repens* | Philippines | Cabanatuan, NE | 1989 | (*5*) | Panicum | PoP | SAMN19488836 |

| | | | | | | | | |
|---|---|---|---|---|---|---|---|---|
| Pd88413 | *Paspalum distichum* (knotgrass) | Philippines | Los Baños, LB | 1988 | (*5*) | Echinochloa | PoEc | SAMN14144143 |
| PtKY18-1 | *Poa trivialis* (rough bluegrass) | USA | Lexington KY | 2018 | This study | Lolium1 | n.d. | SAMN19488838 |
| RrJA49 | *Rhynchelytrum roseum* (natal grass) | Brazil | MG | 2018 | This study | Urochloa2 | PoU2 | SAMN19488839 |
| BP1-FLA | *Setaria faberi* (Japanese bristlegrass) | USA | Eastland, FL | 2018 | This study | Setaria | PoS | SAMN19488840 |
| GFSI1-7-2 | *Setaria italica* (foxtail millet) | Japan | Gifu | 1977 | (*2*) | Setaria | PoS | SAMD00051170 |
| MG05 | *S. italica* | India | Bangalore, KA | 2012 | (*4*) | Setaria | PoS | SAMN04217000 |
| MG08 | *S. italica* | India | Mandya, KA | 2012 | (*4*) | Setaria | PoS | SAMN04217082 |
| U232 | *S. italica* | Uruguay | Minas, LA | 2017 | (*11*) | Setaria | PoS | SAMN19488841 |
| US71 | *Setaria spp.* | USA | Lexington, KY | ND | (*9*) | Setaria | PoS | SAMEA3373385 |
| Arcadia2 | *Setaria viridis* (green foxtail) | USA | Arcadia Park, Lexington, KY | 1998 | (*6*) | Setaria | PoS | SAMN14167122 |
| GRF52 | *S. viridis* | USA | Spindle Top Farm, Lexington, KY | 2001 | (*6*) | Setaria | PoS | SAMN08009556 |

| | | | | | | | | | |
|---|---|---|---|---|---|---|---|---|---|
| KANSV1-4-1 | | *S. viridis* | Japan | Kanagawa | 1975 | (*2*) | Setaria | PoS | SAMD00051178 |
| SA05-144 | | *S. viridis* | Japan | Nagasaki | 2005 | (*2*) | Setaria | PoS | SAMD00051180 |
| SA05-43 | | *S. viridis* | Japan | Nagasaki | 2005 | (*2*) | Setaria | PoS | SAMD00051179 |
| Sv9610 | | *S. viridis* | China | ZJ | 1996 | (*21*) | Setaria | PoS | SAMN04318449 |
| Sv9623 | | *S. viridis* | China | ZJ | 1996 | (*21*) | Setaria | PoS | SAMN04318450 |
| Pg1054 | | *Stenotaphrum secundatum* (St. Augustinegrass) | USA | GA | 1999 | (*11*) | Stenotaphrum | PoSt | SAMN19488842 |
| Pg1204 | | *S. secundatum* | USA | GA | 1999 | This study | Stenotaphrum | PoSt | SAMN19488843 |
| SSFL02-1 | SSFL02 | *S. secundatum* | USA | Disneyworld, FL | 2002 | (*3*) | Stenotaphrum | PoSt | SAMN08009573 |
| SSFL14-3 | | *S. secundatum* | USA | New Smyrna, FL | 2014 | (*6*) | Stenotaphrum | PoSt | SAMN08009574 |
| SSTX16-11 | SSTX16-1 | *S. secundatum* | USA | TX | 2016 | (*11*) | Stenotaphrum | PoSt | SAMN14144144 |
| STAG-MS | | *S. secundatum* | USA | MS | 1980 | This study | Stenotaphrum | PoSt | SAMN19488844 |
| U217 | | *S. secundatum* | Uruguay | Valle Alto, TT | 2015 | (*11*) | Stenotaphrum | PoSt | SAMN19488845 |

| | | | | | | | | | |
|---|---|---|---|---|---|---|---|---|---|
| U233 | | *S. secundatum* | Uruguay | Covidef 1, FL | 2017 | (*11*) | Stenotaphrum | PoSt | SAMN19488846 |
| B2 | | *Triticum aestivum* (wheat) | Bolivia | Quirusillas, SC | 2011 | (*6*) | Triticum1 | n.d. | SAMN05580113 |
| B71 | | *T. aestivum* | Bolivia | Quirusillas, SC | 2012 | (*22*) | Triticum1 | n.d. | SAMN04942725 |
| BdBar | P161 | *T. aestivum* | Bangladesh | Barisal | 2016 | This study | Triticum1 | n.d. | SAMN04940126 |
| BdJes | P162 | *T. aestivum* | Bangladesh | Jessore | 2016 | This study | Triticum1 | n.d. | SAMN04942531 |
| BdKUS | | *T. aestivum* | Bangladesh | Kushtia | 2016 | This study | Triticum1 | n.d. | SAMN14144137 |
| BdMag | | *T. aestivum* | Bangladesh | Magura | 2016 | This study | Triticum1 | n.d. | SAMN14144138 |
| BdMeh | P163 | *T. aestivum* | Bangladesh | Mehepur | 2016 | This study | Triticum1 | n.d. | SAMN04942534 |
| Br2 | | *T. aestivum* | Brazil | PR | 1990 | | Triticum1 | n.d. | |
| Br3 | | *T. aestivum* | Brazil | PR | 1990 | | Triticum1 | n.d. | |
| Br115-7 | Br115.7 | *T. aestivum* | Brazil | PR | 1992 | | Triticum1 | n.d. | |
| Br115-12 | Br115.12 | *T. aestivum* | Brazil | PR | 1992 | | Triticum1 | n.d. | |
| BR116 | Br116.5 | *T. aestivum* | Brazil | PR | 1992 | (*10*) | Triticum1 | n.d. | SAMN14144139 |
| BR118 | | *T. aestivum* | Brazil | PR | 1992 | (*10*) | Triticum1 | n.d. | SAMN14144140 |
| Br126-1 | Br126.1 | *T. aestivum* | Brazil | PR | 1992 | | Triticum1 | n.d. | |

| | | | | | | | | | |
|---|---|---|---|---|---|---|---|---|---|
| Br127-1 | Br127.1 | *T. aestivum* | Brazil | PR | 1992 | | Triticum1 | n.d. | |
| Br127-11 | Br127.11 | *T. aestivum* | Brazil | PR | 1992 | | Triticum1 | n.d. | |
| Br130 | | *T. aestivum* | Brazil | MS | 1990 | (*6*) | Triticum1 | n.d. | SAMN08009547 |
| Br130-8 | Br130.8 | *T. aestivum* | Brazil | PR | 1992 | | Triticum1 | n.d. | |
| Br130-9 | Br130.9 | *T. aestivum* | Brazil | PR | 1992 | | Triticum1 | n.d. | |
| BR32 | BR0032 | *T. aestivum* | Brazil | - | 1991 | (*9*) | Triticum1 | n.d. | SAMEA4708260 |
| Br46 | | *T. aestivum* | Brazil | MS | 1990 | | Triticum1 | n.d. | |
| Br48 | | *T. aestivum* | Brazil | MS | 1990 | (*2*) | Triticum1 | n.d. | SAMD00084261 |
| Br49 | | *T. aestivum* | Brazil | MS | 1990 1990 | | Triticum1 | n.d. | |
| Br7 | | *T. aestivum* | Brazil | PR | 1990 | (*6*) | Triticum1 | n.d. | SAMN08009545 |
| Br8 | | *T. aestivum* | Brazil | PR | 1992 | | Triticum1 | n.d. | |
| Br80 | | *T. aestivum* | Brazil | - | 1991 | (*6*) | Triticum1 | n.d. | SAMN08009546 |
| BR81 | | *T. aestivum* | Brazil | - | 1991 | (*23*) | Triticum1 | n.d. | SAMN14144142 |
| BTGP-1b | | *T. aestivum* | Bangladesh | Mehepur | 2017 | (*14*) | Triticum1 | n.d. | SAMN19488847 |
| BTGP-6e | | *T. aestivum* | Bangladesh | Mehepur | 2017 | (*14*) | Triticum1 | n.d. | SAMN19488848 |
| PY0925 | | *T. aestivum* | Brazil | Perdizes, SP | 2009 | (*9*) | Triticum1 | n.d. | SAMEA4029894 |
| Py221 | Py22.1 | *T. aestivum* | Brazil | PR | 2007 | (*3*) | Triticum1 | n.d. | SAMN05725179 |
| PY36 | PY36.1 | *T. aestivum* | Brazil | Brasilia, DF | 2007 | (*11*) | Triticum1 | n.d. | SAMEA4029897 |
| PY5003 | | *T. aestivum* | Brazil | Londrina, PR | 2005 | (*11*) | Triticum1 | n.d. | SAMEA4029888 |
| PY5010 | | *T. aestivum* | Brazil | Londrina, PR | 2005 | (*9*) | Lolium1 | n.d. | SAMEA4029898 |
| Py5020 | | *T. aestivum* | Brazil | Londrina, PR | 2005 | (*3*) | Triticum1 | n.d. | SAMN05762829 |
| PY5033 | PY05033 | *T. aestivum* | Brazil | Londrina, PR | 2005 | (*9*) | Triticum1 | n.d. | SAMEA4029889 |
| PY6017 | PY06017 | *T. aestivum* | Brazil | Coromandel, MG | 2006 | (*9*) | Triticum1 | n.d. | SAMEA4029890 |

| | | | | | | | | | |
|---|---|---|---|---|---|---|---|---|---|
| PY6025 | | *T. aestivum* | Brazil | MG | 2006 | (*8*) | Triticum1 | n.d. | SAMEA4029891 |
| PY6045 | | *T. aestivum* | Brazil | GO | 2006 | (*8*) | Triticum1 | n.d. | SAMEA4029900 |
| PY86 | PY86.1 | *T. aestivum* | Brazil | PR | 2008 | (*8*) | Lolium1 | n.d. | SAMEA4029893 |
| T12-8 | | *T. aestivum* | Brazil | Floresta, PR | 1988 | This study | Triticum1 | n.d. | SAMN19488849 |
| T13-3 | | *T. aestivum* | Brazil | Floresta, PR | 1988 | This study | Triticum1 | n.d. | SAMN19488850 |
| T2-1 | T-0002 | *T. aestivum* | Brazil | Londrina, PR | 1987 | This study | Triticum1 | n.d. | SAMN19488852 |
| T21-1 | | *T. aestivum* | Brazil | Floresta, PR | 1988 | This study | Triticum1 | n.d. | SAMN19488853 |
| T25 | | *T. aestivum* | Brazil | São Jorge do Ivaí, PR | 1988 | This study | Triticum1 | n.d. | SAMN08009575 |
| T3-1 | | *T. aestivum* | Brazil | Vicentinópolis, GO | 1986 | This study | Triticum1 | n.d. | SAMN19488854 |
| T30-2 | | *T. aestivum* | Brazil | PR | 1989 | This study | Triticum1 | n.d. | SAMN19488855 |
| T37-2 | | *T. aestivum* | Brazil | PR | 1989 | This study | Triticum1 | n.d. | SAMN19488856 |
| T4-2 | | *T. aestivum* | Brazil | Floresta, PR | 1988 | This study | Triticum1 | n.d. | SAMN19488857 |
| T42-2 | | *T. aestivum* | Brazil | PR | 1989 | This study | Triticum1 | n.d. | SAMN19488858 |

| | | | | | | | | | |
|---|---|---|---|---|---|---|---|---|---|
| T46-2 | | *T. aestivum* | Brazil | PR | 1989 | This study | Triticum1 | n.d. | SAMN19488859 |
| T5-3 | | *T. aestivum* | Brazil | Palotina, PR | 1988 | This study | Triticum1 | n.d. | SAMN19488861 |
| T50-3 | T-0050 | *T. aestivum* | Brazil | MG | 1989 | This study | Triticum1 | n.d. | SAMN19488862 |
| WB032i | 12.1.032i | *T. aestivum* | Brazil | PR | 2012 | (*1*) | Triticum1 | n.d. | SAMN19488863 |
| WB053i | 12.1.053i | *T. aestivum* | Brazil | SP | 2012 | (*1*) | Triticum1 | n.d. | SAMN19488864 |
| WB127 | 12.1.127 | *T. aestivum* | Brazil | MA | 2012 | (*1*) | Triticum1 | n.d. | SAMN19488865 |
| WB169 | 12.1.169 | *T. aestivum* | Brazil | MA | 2012 | (*1*) | Triticum1 | n.d. | SAMN19488866 |
| WB181 | 12.1.181 | *T. aestivum* | Brazil | RS | 2012 | (*1*) | Triticum1 | n.d. | SAMN18576982 |
| WB205 | 12.1.205 | *T. aestivum* | Brazil | RN | 2012 | (*1*) | Triticum1 | n.d. | SAMN18576983 |
| WB217 | 12.1.217 | *T. aestivum* | Brazil | RS | 2012 | (*1*) | Triticum1 | n.d. | SAMN18576984 |
| WB225 | 12.1.225 | *T. aestivum* | Brazil | RS | 2012 | (*1*) | Triticum1 | n.d. | SAMN18576985 |
| WB241 | 12.1.241 | *T. aestivum* | Brazil | MS | 2012 | (*1*) | Triticum1 | n.d. | SAMN18576987 |
| WB291 | 12.1.291 | *T. aestivum* | Brazil | PR | 2012 | (*1*) | Triticum1 | n.d. | SAMN18576988 |
| WB37 | 12.1.037 | *T. aestivum* | Brazil | GO | 2012 | (*1*) | Triticum1 | n.d. | SAMN18576980 |
| WBKY11 | WBKY11-15 | *T. aestivum* | USA | Lexington, KY | 2011 | (*6*) | Lolium1 | n.d. | SAMN08009578 |
| WBSS | | *T. aestivum* | Brazil | - | - | (*6*) | Triticum1 | n.d. | SAMN08009579 |
| WHTQ | | *T. aestivum* | Brazil | - | - | (*6*) | Triticum1 | n.d. | SAMN08009580 |
| P3 | | *Triticum durum* (durum wheat) | Paraguay | CY | 2012 | (*3*) | Triticum1 | n.d. | SAMN08009568 |

| | | | | | | | | | |
|---|---|---|---|---|---|---|---|---|---|
| Ub007i | 12.0.007i | *Urochloa brizantha* (palisade grass) | Brazil | PR | 2012 | (*1*) | Triticum1 | n.d. | SAMN07829570 |
| Ub009i | 12.0.009i | *U. brizantha* | Brazil | PR | 2012 | (*1*) | Triticum1 | n.d. | SAMN07829571 |
| Ub012i | 12.0.012i | *U. brizantha* | Brazil | PR | 2012 | (*1*) | Triticum1 | n.d. | SAMN07829572 |
| UbJA112 | | *U. brizantha* | Brazil | MG | 2018 | This study | Stenotaphrum | PoSt | SAMN19488867 |
| UbJA92 | | *U. brizantha* | Brazil | MG | 2018 | This study | Urochloa2 | PoU2 | SAMN19488868 |
| Ud8401 | Bd8401 | *Urochloa distachya* (tropical signalgrass) | Philippines | - | 1984 | (*6*) | Urochloa2 | PoP | SAMN08009543 |
| Um8309 | Bm8309 | *Urochloa mutica* (buffalo grass) | Philippines | Los Baños, LB | 1983 | (*5*) | Urochloa1 | PoU1 | SAMN19488868 |
| Um88324 | Bm88324 | *U. mutica* | Philippines | Cabanatuan, NE | 1988 | (*5*) | Urochloa1 | PoU1 | SAMN08009544 |
| Um8946 | Bm8946 | *U. mutica* | Philippines | Imus, Cv | 1989 | (*5*) | Urochloa2 | PoU1 | SAMN19488870 |
| Up35 | Br35 | *Urochloa plantaginea* (creeping signalgrass) | Brazil | PR | 1990 | (*10*) | Urochloa3 | PoSt | SAMN14144141 |
| P25 | P-0025 | *Urochloa spp.* | Paraguay | CY | 2014 | (*11*) | Lolium1 | n.d. | SAMN19488822 |

Table S2. Haplotype designations of *Lolium*/*Triticum* lineage isolates

| Haplotype[A] | Isolates | Year of [B] isolation | MATallele | Mitotype[D] | PWT3 allele[E] | PWT3 source |
|---|---|---|---|---|---|---|
| *Lolium* | | | | | | |
| Founder, 1 | ATCC64557 | 1980 | 1-2 | PoE1 | A | PoU1 |
| 2 | Br58*,[F] Ds555i*, As073* | 1990 | 1-2 | PoX | A | PoU1 |
| 3 | CHRF, CHW, FH, GG11, HO, LpKY, PgKY, PgPA, TP2 | 1996 | 1-2 | PoX | A | PoU1 |
| 4 | PL2-1, PtKY18-1* | 2002 | 1-2 | PoX | A | PoU1 |
| 5 | PL3-1, WBKY11* | 2002 | 1-1 | PoX | A/Atp[G] | PoU1 |
| 6 | Po221, PY5010*, TF05-1, TF15-1 | 2005 | 1-2 | PoX | A | PoU1 |
| 7 | PY86* | 2008 | 1-1 | PoX | A | PoU1 |
| 8 | Pg1213-22 | 2010 | 1-1 | PoX | A | PoU1 |
| 9 | P25* | 2012 | 1-2 | PoX | A | PoU1 |
| 10 | P28* | 2012 | 1-2 | PoX | A | PoU1 |
| 11 | As321*, As345*, As347*, Ecrus326*, Ub368* | 2012 | 1-2 | n.d. | A | PoU1 |
| 12 | WB217* | 2012 | 1-2 | n.d. | B | PoSt |
| 13 | U234 | 2017 | 1-1 | PoX | A | PoU1 |
| *Triticum* | | | | | | |
| 1 | T3-1, T47-3 (*Triticum* founder lineage) | 1985 | 1-1 | PoX | A | PoU1 |
| 2 | T1-1, T5-3, T12-8, BR32, BR81 | 1987 | 1-1 | PoX | Atm | de novo |

| | | | | | | |
|---|---|---|---|---|---|---|
| 3 | T2-1 | 1987 | 1-1 | PoX | Atm | de novo |
| 4 | T21-1, WB127, WB169 | 1988 | 1-2 | PoX | Atc | 2PoE1 |
| 5 | T7-3, T30-2, T42-2 | 1988 | 1-1 | PoX | A | PoU1 |
| 6 | Br8, T13-3 | 1988 | 1-1 | PoX | B | PoSt |
| 7 | T37-2 | 1989 | 1-1 | PoX | A | PoU1 |
| 8 | T50-3 | 1989 | 1-1 | PoX | B | PoSt |
| 9 | Br2, Br3, Br48, T4-2, T25 | 1988 | 1-1 | PoX | B | PoSt |
| 10 | Br7 | 1990 | 1-2 | PoX | B | PoSt |
| 11 | Br46 | 1990 | 1-1 | n.d. | Atm | de novo |
| 12 | Br49 | 1990 | 1-1 | n.d. | Atm | de novo |
| 13 | BR80 | 1991 | 1-1 | PoX | A | PoE1 |
| 14 | Py221, PY6025, PY6045, Ub007i*, Ub009i*, Ub012i*, WBSS | 1992 | 1-1 | PoX | Atc | 2PoE1 |
| 15 | BR116, T46-2 | 1989 | 1-2 | PoX | Atm | de novo |
| 16 | PY6017, WHTQ | 1992 | 1-1 | PoX | Atc | 2PoE1 |
| 17 | BR118 | 1992 | 1-1 | PoX | A | PoU1 |
| 18 | BR130 | 1992 | 1-1 | PoX | A**H | PoU1 |
| 19 | Br108-1 | 1992 | 1-1 | n.d. | Atm | de novo |
| 20 | Br115-7 | 1992 | 1-1 | n.d. | A | PoU1 |
| 21 | Br127-11 | 1992 | 1-1 | n.d. | Atm | de novo |
| 28, 22 | Br130-8 | 1992 | 1-1 | n.d. | Atm | de novo |

| | | | | | | |
|---|---|---|---|---|---|---|
| 23 | Br130-9 | 1992 | 1-2 | n.d. | Atc | PoE1 |
| 24 | Br127-1 | 1992 | 1-1 | n.d. | A | PoU1 |
| 25 | Br115-12 | 1992 | 1-1 | n.d. | A | PoU1 |
| 26 | Br126-1 | 1992 | 1-1 | n.d. | A | PoU1 |
| 27 | B2, P3, PY5003 | 2005 | 1-2 | PoX | Atm | de novo |
| 28 | PY5033 | 2005 | 1-1 | PoX | Atc | 2PoE1 |
| 29 | PY5020, WB014 | 2005 | 1-1 | PoX | Atc | 2PoE1 |
| 30 | PY36 | 2007 | 1-1 | PoX | Atc | 2PoE1 |
| 31 | Ce642i*, Ce535*, Ei534i*, EiJA22*, Mr051i, PY0925, WB37, WB053i, | 2009 | 1-1 | PoX | Atc | 2PoE1 |
| 32 | B71, BdBar, BdJes, BdKUS, BdMag, BdMeh, BTBa-B1, BTBa-B2, BTGP-1b, BTGP-6e, WB181 | 2012 | 1-1 | PoSt | Atc | 2PoE1 |
| 33 | WB078 | 2012 | | n.d. | Atc | 2PoE1 |
| 34 | P29*, WB032i, WB205, WB225, WB241, WB291, Ecan194* | 2012 | 1-1 | PoSt | Atc | 2PoE1 |

[A] Haplotypes were assigned based on pairwise SNP divergence. Isolates with < 100 SNPs/Mb difference were assumed to have the same chromosomal haplotype.

[B] Date that the first isolate representing the haplotype was collected

[C] Mating type determined by interrogating genome sequence

[D] Mitochondrial genotype

[E] Allele designations from reference (*1*)

[F] * indicates strain was isolated from a different host species (i.e. non-wheat/*Lolium*)

[G] WBKY11 contains a new transposons insertion in the promoter of the *PWT3* gene

[H] ** indicates a variant allele

Table S3. PoT/PoL haplotypes harboring reciprocal crossover products

| Chromosome | pre-X over SNP pos. | pre-X over alleles | post-X-over SNP pos. | post-X-over alleles | Haplotype(s) |
|---|---|---|---|---|---|
| **1** | 6278974 | E | 6280676 | St | PoT26 |
| | 6278974 | St | 6280676 | E | PoT2 |
| **3** | **7340282** | **St** | **7354813** | **E** | **PoL6, PoL7** |
| | **7340282** | **E** | **7354813** | **St** | **PoT14, PoT31, PoT4, PoT30, PoT20** |
| **4** | 164223 | E | 182460 | St | PoT5, PoT22, PoT12 |
| | 164223 | St | 182460 | E | PoT34 |
| | 357691 | U1 | 385442 | St | PoT13 |
| | 357691 | St | 385442 | U1 | PoT31, PoT6 |
| | 4471461 | Lu | 4573068 | E | PoL3, PoL4, PoL7, PoL13 |
| | 4471461 | E | 4573068 | Lu | PoL5, PoL8 |
| **6** | 4440024 | X | 4441248 | E | PoT2 |
| | 4440024 | E | 4441248 | X | PoT18 |
| | 6021357 | X | 6057434 | St | PoT10, PoT17, PoT21 |
| | 6021357 | St | 6057434 | X | PoT14, PoT4, PoT30, PoT20, PoT24 |
| **7** | 385442 | St | 525372 | X | PoT9, PoT3 |
| | 385442 | X | 525372 | St | PoT32, PoT31, PoT2, PoT34, PoT13, PoT18 |
| | 3517584 | St | 3538663 | E | PoT14, PoT34, PoT30 |
| | 3517584 | E | 3538663 | St | PoT5, PoT17, PoT7, PoT24, PoT12 |

Table S4. Donor genome proportions inherited by each PoL/PoT haplotype.

| Haplotype | PoE1+PoE2 | PoLu | PoSt | PoU1 | PoX |
|---|---|---|---|---|---|
| PoL1 | 65.9 | 0.3 | 0.3 | 33.3 | 0.2 |
| PoL2 | 58.5 | 5.9 | 2.6 | 16.9 | 16.0 |
| PoL3 | 55.6 | 4.9 | 2.4 | 17.8 | 19.3 |
| PoL4 | 55.6 | 4.9 | 2.5 | 17.7 | 19.3 |
| PoL5 | 55.4 | 2.7 | 2.2 | 18.9 | 20.8 |
| PoL6 | 54.2 | 5.1 | 3.3 | 15.2 | 22.2 |
| PoL7 | 56.3 | 5.2 | 2.9 | 16.5 | 19.2 |
| PoL8 | 55.5 | 2.6 | 2.1 | 18.9 | 21.0 |
| PoL9 | 52.9 | 6.0 | 4.8 | 17.2 | 19.2 |
| PoL10 | 57.9 | 0.9 | 6.7 | 19.9 | 14.6 |
| PoL11 | 53.4 | 6.0 | 11.9 | 15.0 | 13.7 |
| PoL13 | 53.2 | 5.1 | 10.1 | 16.1 | 15.5 |
| PoT1 | 52.8 | 0.4 | 0.7 | 19.2 | 26.9 |
| PoT2 | 47.8 | 1.2 | 23.7 | 11.4 | 15.9 |
| PoT3 | 36.0 | 0.7 | 31.9 | 18.9 | 12.5 |
| PoT4 | 48.0 | 7.9 | 16.8 | 9.7 | 17.6 |
| PoT5 | 47.1 | 1.7 | 26.3 | 12.4 | 12.5 |
| PoT6 | 37.3 | 0.9 | 33.8 | 12.1 | 15.9 |
| PoT7 | 50.2 | 1.0 | 18.0 | 12.4 | 18.3 |
| PoT8 | 42.4 | 0.6 | 27.4 | 10.0 | 19.5 |
| PoT9 | 42.6 | 1.0 | 26.7 | 13.8 | 16.0 |
| PoT10 | 45.6 | 2.1 | 24.6 | 11.2 | 16.5 |
| PoT11 | 47.0 | 0.6 | 21.6 | 14.9 | 15.8 |
| PoT12 | 45.7 | 1.1 | 21.7 | 12.1 | 19.4 |
| PoT13 | 49.6 | 0.8 | 20.2 | 10.5 | 18.9 |
| PoT14 | 48.1 | 1.2 | 20.0 | 14.1 | 16.6 |
| PoT15 | 41.1 | 1.4 | 31.5 | 7.5 | 18.4 |
| PoT16 | 50.6 | 0.9 | 19.7 | 12.3 | 16.5 |
| PoT17 | 48.6 | 1.8 | 16.5 | 16.6 | 16.5 |
| PoT18 | 50.5 | 1.1 | 13.8 | 16.3 | 18.2 |

| | | | | | |
|---|---|---|---|---|---|
| PoT19 | 49.1 | 0.7 | 14.2 | 19.2 | 16.8 |
| PoT20 | 47.7 | 1.0 | 19.4 | 16.1 | 15.8 |
| PoT21 | 46.5 | 1.6 | 18.8 | 15.1 | 18.0 |
| PoT22 | 47.2 | 1.8 | 19.1 | 16.0 | 15.9 |
| PoT24 | 48.0 | 1.3 | 18.3 | 16.7 | 15.7 |
| PoT25 | 40.5 | 1.3 | 27.4 | 11.0 | 19.8 |
| PoT26 | 43.8 | 1.6 | 24.2 | 10.9 | 19.6 |
| PoT27 | 43.3 | 0.9 | 29.9 | 6.6 | 19.4 |
| PoT28 | 50.1 | 1.1 | 21.3 | 11.0 | 16.4 |
| PoT30 | 46.5 | 0.7 | 26.4 | 11.3 | 15.1 |
| PoT31 | 55.5 | 1.2 | 16.8 | 12.2 | 14.2 |
| PoT32 | 57.1 | 0.6 | 16.3 | 12.6 | 13.5 |
| PoT34 | 50.1 | 1.7 | 22.4 | 11.2 | 14.6 |

Table S5. PoLu gene conversions on chromosome 4

| Start position | End position | length (bp) |
|---|---|---|
| 897,550 | 899,300 | 1,750 |
| 2,338,030 | 2,363,600 | 25,570 |
| 2,527,700 | 2,541,200 | 13,500 |
| 3,164,000 | 3,169,000 | 5,000 |
| 3,205,500 | 3,233,000 | 27,500 |
| 5,058,000 | 5,072,300 | 14,300 |

Table S6. Proportions of respective donor populations present in PoT/PoL lineages.

| | PoE1+PoE2 | PoLu | PoSt | PoU1 | PoX |
|---|---|---|---|---|---|
| Chromosome equivalents | | | | | |
| Chr1 | 0.81 | 0.01 | 0.43 | 0.21 | 0.22 |
| Chr2 | 0.50 | 0.05 | 0.75 | 0.57 | 0.64 |
| Chr3 | 1.00 | 0.02 | 0.88 | 0.00 | 0.29 |
| Chr4 | 0.17 | 0.57 | 1.00 | 0.86 | 0.49 |
| Chr5 | 0.76 | 0.07 | 0.58 | 0.39 | 0.27 |
| Chr6 | 1.00 | 0.06 | 0.78 | 0.01 | 0.57 |
| Chr7 | 0.66 | 0.11 | 1.00 | 0.61 | 0.36 |
| Genome equivalents | | | | | |
| PoL | 0.62 | 0.08 | 0.2 | 0.38 | 0.34 |
| PoT | 0.6 | 0.11 | 0.74 | 0.24 | 0.28 |
| PoT/PoL | 0.7 | 0.13 | 0.77 | 0.38 | 0.41 |

Table S7. Changes in the frequencies of various *PWT3* alleles over time

| Allele | 1985 (1)[A] | 1986 (2) | 1987 (3) | 1988 (7) | 1989 (7) | 1990-2000 (22) | 2001-2010 (9) | 2011-2020 (27)[B] |
|---|---|---|---|---|---|---|---|---|
| A | 100 | 100 | | 14.3 | 57.1 | 31.8 | 0.0 | 6.9 |
| B | | | 100 | 42.9 | 28.6 | 22.7 | 0.0 | 3.4 |
| Atc | | | | 14.3 | 0.0 | 4.5 | 88.9 | 79.3 |
| Atm | | | | 28.6 | 14.3 | 36.4 | 11.1 | 6.9 |
| Atp | - | - | - | - | - | - | - | 3.4 |

[A] Values in parentheses are the numbers of isolates surveyed from each collection period.

[B] Isolates from Bangladesh and India were not included in this numbers because they originated as clones of a single haplotype.

**Supplementary Figure Legends**

Figure S1. Relaxed host specificity in the PoT/PoL lineages (see Figure 1C). Shown is a neighboring joining tree for the PoT and PoL lineages, based on pairwise distance data. Colored circles indicate the different host genera from which particular haplotypes were isolated. Note that certain haplotypes were capable of infecting multiple host grasses.

Figure S2. Discordant phylogenies for two linked loci on chromosome 7. Phylogenetic trees were generated for 20 kb of sequence surrounding markers MPG1 (Chr7:159,716-179,715) and CH7BAC7 (Chr7:1,173,687-1,193,686) using RAxML version 8 (Stamatakis, 2014), assuming a GTR gamma model and with 100 bootstrap replications. Tips were re-ordered and plotted using the cophylo function in phytools (Revell, 2012). PoT lineage members are labeled in blue and PoL members in purple.

Figure S3. DAPC analysis of *P. oryzae* populations. A) Use of Bayesian Information Criterion to determine most probable number of discrete populations. Max clusters was set at K=30 and the find.clusters function was performed on the first 80 principle components, and was repeated for 1 million iterations. The lowest mean BIC value was at K=16. Clustering and plotting was implemented using the poppr package (Kamvar et al. 2014). B) Assignment of population memberships using discriminant analysis of principle components (DAPC). The first 80 principle components were retained and used for discriminant analysis of population memberships at K values from 10 to 25.

Figure S4. Neighbor network showing extensive reticulation within the PoT and PoL clades. Pairwise distance data were used to build a neighbor network using the Splits Network Algorithm in SplitsTree (*1, 2*). Lineages identified using DAPC are shown in shaded "clouds." Isolates not mapping close to their assigned groups are given individual strain labels. Scale indicates sequence divergence and the proportion of distances represented by edges (fit) is shown.

Figure S5. The PoLu lineage is admixed with contributions from PoU1, PoSt and an unknown donor. Shown are haplotype divergence plots for *Luziola* pathogens U168 and U171, and the *Hakonechloa* pathogen FPH-2015-44. Plots lines are included if the candidate donor lineage showed low haplotype divergence to at least one chromosome region in the test strain. Numbers of comparator isolates in each lineage were PoSt (n = 10), PoU1 (n= 3) , and PoU3 (a sub-population of PoSt, n = 2,). U168 and U171 are essentially clones of one another.

Figure S6.Chromopaintings of the PoL/PoT haplotypes annotated to show informative features. Plots were generated from the same dataset used for Fig. 4. Segments predicted to have been donated via sib-mating are highlighted with dotted lines, those that appear to have come from backcrosses are encompassed by areas with rounded corners, and sequences from secondary introgressions have borders with square corners. Representative reciprocal crossovers are marked with botted line arrowheads. The locations of the *PWT3* and $PWT6^{null}$ host-specificity loci, the PoT-diagnostic MoT3 and C17 markers, as well as the phylogenetically informative markers MPG1, CH7BAC7, are marked with triangles. MoT3 maps at a locus that was not painted due to ChromoPainter being unable to analyze sites that are missing/repeated in some strains. Regions used for tip dating are indicated with white boxes and those analyzed for secondary introgressions are marked with black boxes. Asterisks show that $pwt3^{B}$ reside on a PoSt introgression segment.

Figure S7 Identification of secondary introgressions from the PoSt and PoE1 lineages. A) Maximum Likelihood Tree built using binary haplotype data for the PoSt sequences on chromosome 7 between positions 171523 and 177915. Bootstrap values are shown. B) ML tree for haplotype data for PoE1 sequences on chromosome 1 between positions 4741547 and 4839703. Both trees were built using the bingamma substitution model and 100 bootstrap replications. C) Haplotypes for select PoT isolates and candidate donors on chromosome 7 between positions 171,523 to 177,030. Points are plotted at positions where each isolate has a

SNP relative to the B71 reference. PoT tracks are shown adjacent to donor having identical haplotypes. Variant donor haplotypes are also shown. Tracks are colored according to donor (see legend for A and B). D). Haplotypes for representative PoT isolates and candidate donors on chromosome 1 between positions 4741547 and 4839703.

Figure S8. Null/non-functional alleles of the *PWT6* and *PWT3* host-specificity genes were contributed to the swarm via admixture. A) Drawn at the top is a schematic of a 250 kb region spanning the *PWT6* gene in the PoE1 isolate CD156. Colored boxes show retrotransposon insertions, with flanking arrowheads marking LTRs. Solo-LTRs are represented with lone arrowheads. Red lines mark 5S rRNA genes. Strains ATCC64557 (PoL1), LpKY97 (PoL2), and B71 (PoT32) all inherited a $pwt6^{null}$ allele that was contributed to the swarm by the PoU1 donor and they have the same locus structure as isolate, Um88324 (PoU1). Strain T4-2 (PoT9) inherited a slight variant of the null allele that was contributed by the PoSt donor and its locus structure matches that of PoSt isolate, Up35. B) Zoomed in region showing the synteny breakpoints in a 5S rRNA gene. C) RIP patterns expected if MGL transposition in PoT1. i) Most MGL copies in the PoT1 genome should have unique sequences due to past RIP events. Transposition of MGL into $PWT3^{A}$ produces a new copy identical to the source element (asterisked). ii) The new copy would accumulate RIP mutations in subsequent admixture matings - recognizable as transitions relative to the source element - uni-directional after one mating (note: source copy is shown in square brackets due to the possibility of segregation), and bidirectional after two or more (iii). D) RIP patterns expected if the MGL insertion originated in a PoSt introgression segment. i) Source element will likely be RIP'd in the first admixture mating (ii). RIP mutations could be uni-directional or bi-directional, depending on transposition timing and number of subsequent matings post admixture (iii). E) BLAST trace-back operations strings showing nucleotide differences between the MGL insertion in $PWT3^{Atm}$ and the most similar copies in Up35 (PoSt) and F) T3-1 (PoT1).

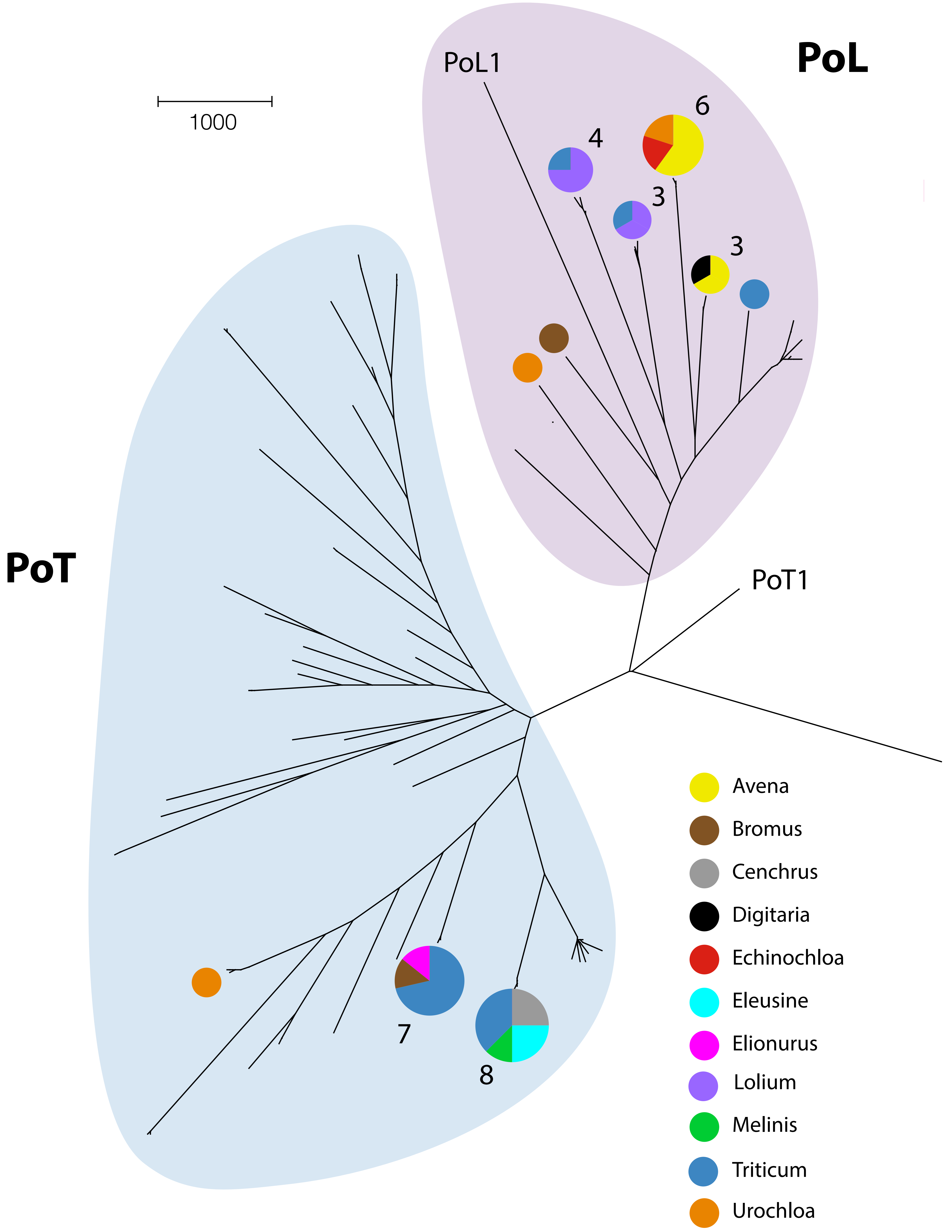
PoL
PoL1
1000
6
4
3
3
PoT
PoT1
7
8
Avena
Bromus
Cenchrus
Digitaria
Echinochloa
Eleusine
Elionurus
Lolium
Melinis
Triticum
Urochloa

**Figure S1**

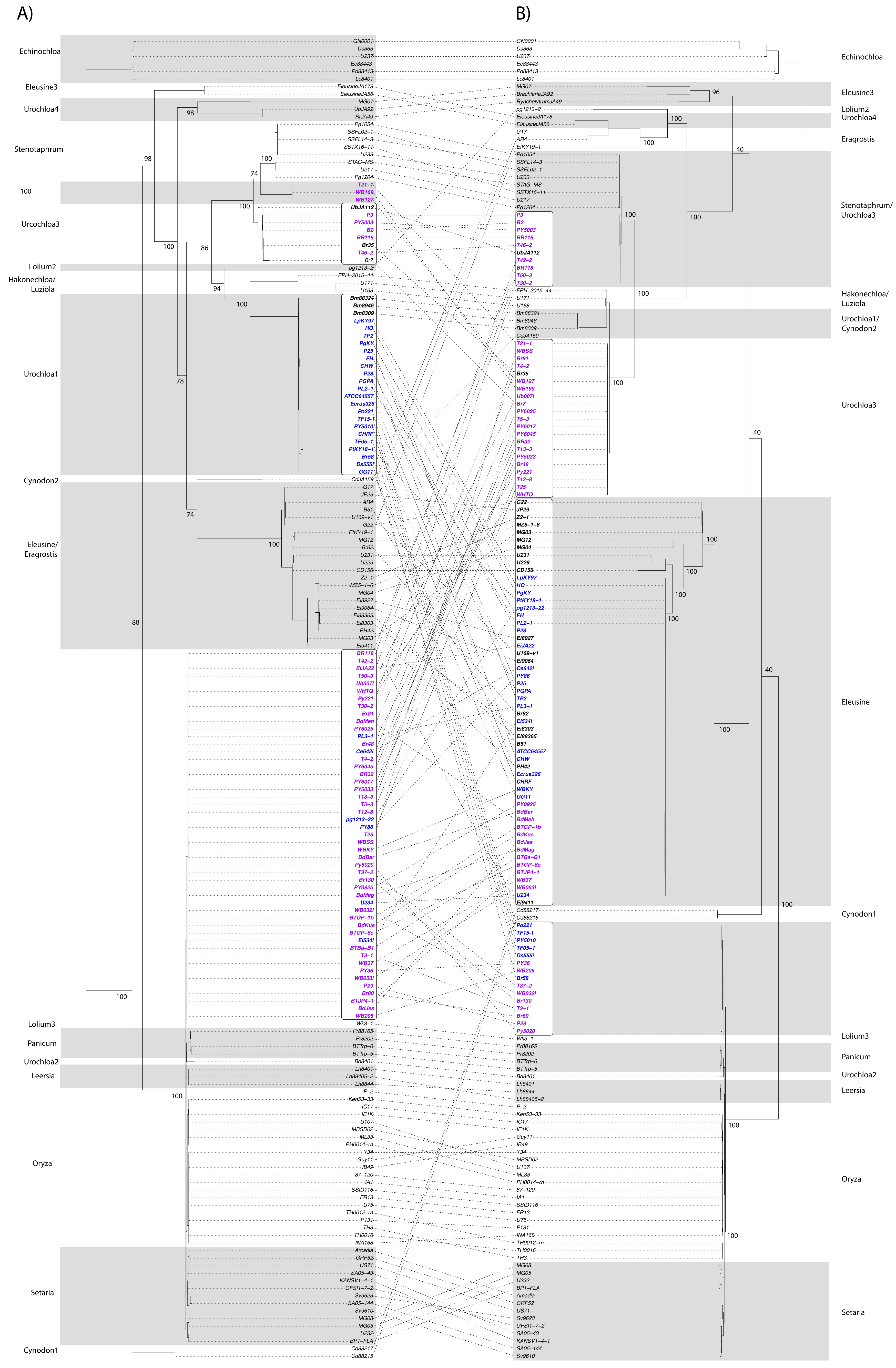
A)
B)
Echinochloa
Eleusine3
Urochloa4
Stenotaphrum
Urcochloa3
Lolium2
Hakonechloa/ Luziola
Urochloa1
Cynodon2
Eleusine/ Eragrostis
Lolium3
Panicum
Urochloa2
Leersia
Oryza
Setaria
Cynodon1
Lolium2 Urochloa4
Eragrostis
Stenotaphrum/ Urochloa3
Urochloa1/ Cynodon2
Eleusine

Figure S2

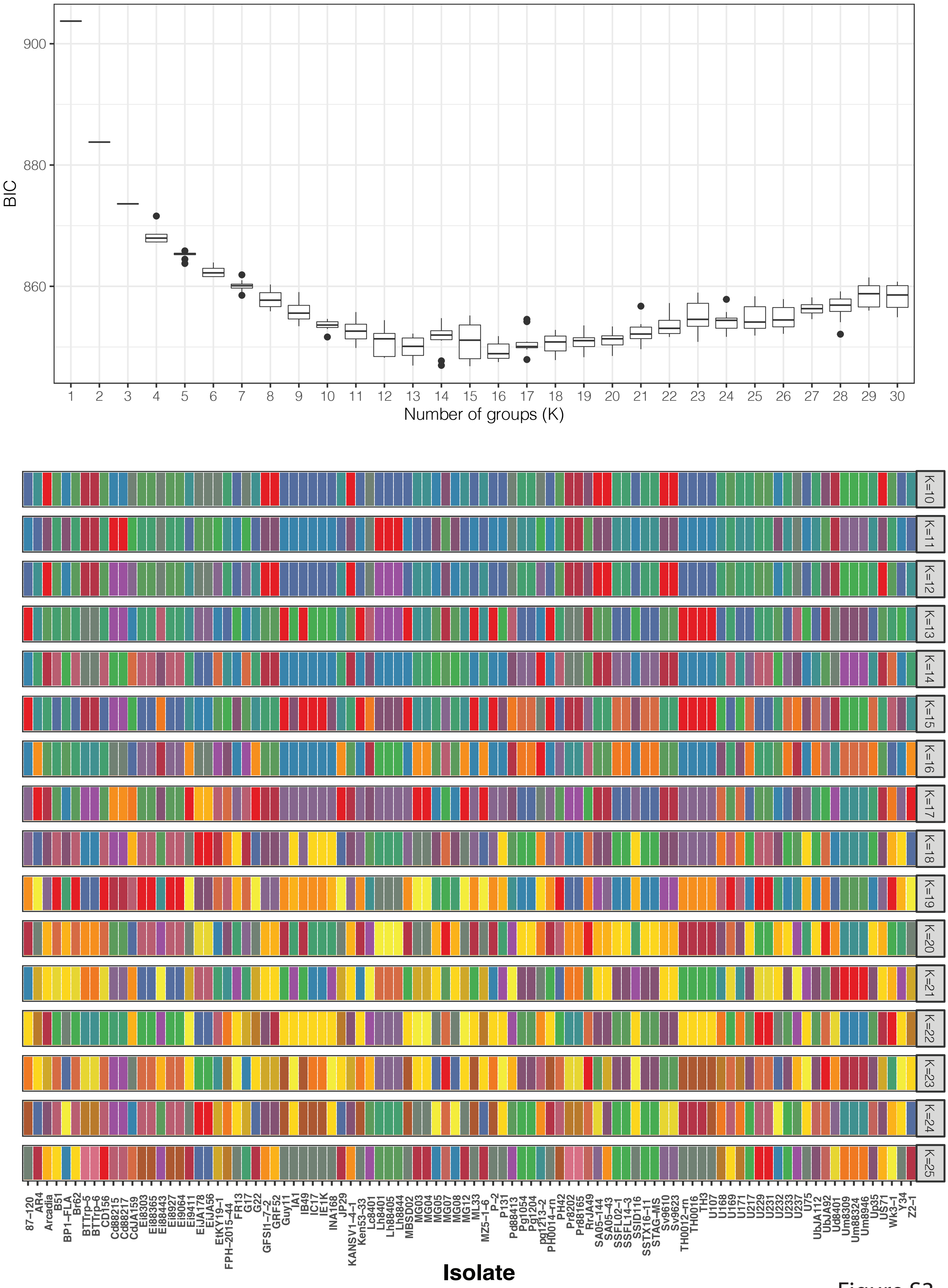

BIC
900
880
860
Number of groups (K)
K=10
K=11
K=12
K=13
K=14
K=15
K=16
K=17
K=18
K=19
K=20
K=21
K=22
K=23
K=24
K=25
87–120
AR4
Arcadia
B51
BP1–FLA
Br62
BTTrp–5
BTTrp–6
CD156
Cd88215
Cd88217
CdJA159
Ei8303
Ei88365
Ei88443
Ei8927
Ei9064
Ei9411
EiJA178
EiJA56
EtKY19–1
FPH–2015–44
FR13
G17
G22
GFSI1–7–2
GRF52
Guy11
IA1
IB49
IC17
IE1K
INA168
JP29
KANSV1–4–1
Ken53–33
Lc8401
Lh8401
Lh88405
Lh8844
MBSD02
MG03
MG04
MG05
MG07
MG08
MG12
ML33
MZ5–1–6
P–2
P131
Pd88413
Pg1054
Pg1204
pg1213–2
PH0014–rn
PH42
Pr8202
Pr88165
RrJA49
SA05–144
SA05–43
SSFL02–1
SSFL14–3
SSID116
SSTX16–11
STAG–MS
Sv9610
Sv9623
TH0012–rn
TH0016
TH3
U107
U168
U169
U171
U217
U229
U231
U232
U233
U237
U75
UbJA112
UbJA92
Ud8401
Um8309
Um88324
Um8946
Up35
US71
Wk3–1
Y34
Z2–1
Isolate

Figure S3

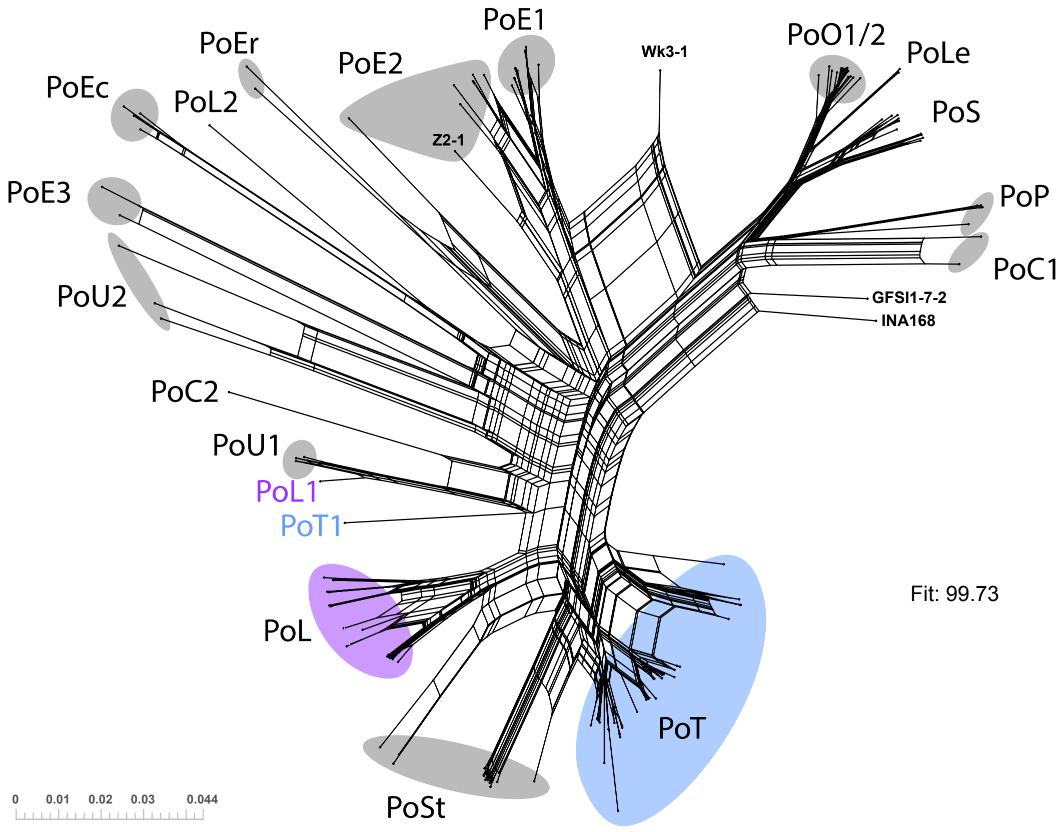
PoE1
PoEr
PoE2
Wk3-1
PoO1/2
PoLe
PoEc
PoL2
PoS
Z2-1
PoE3
PoP
PoC1
PoU2
GFSI1-7-2
INA168
PoC2
PoU1
PoL1
PoT1
Fit: 99.73
PoL
PoT
PoSt
0
0.01
0.02
0.03
0.044

**Figure S4**

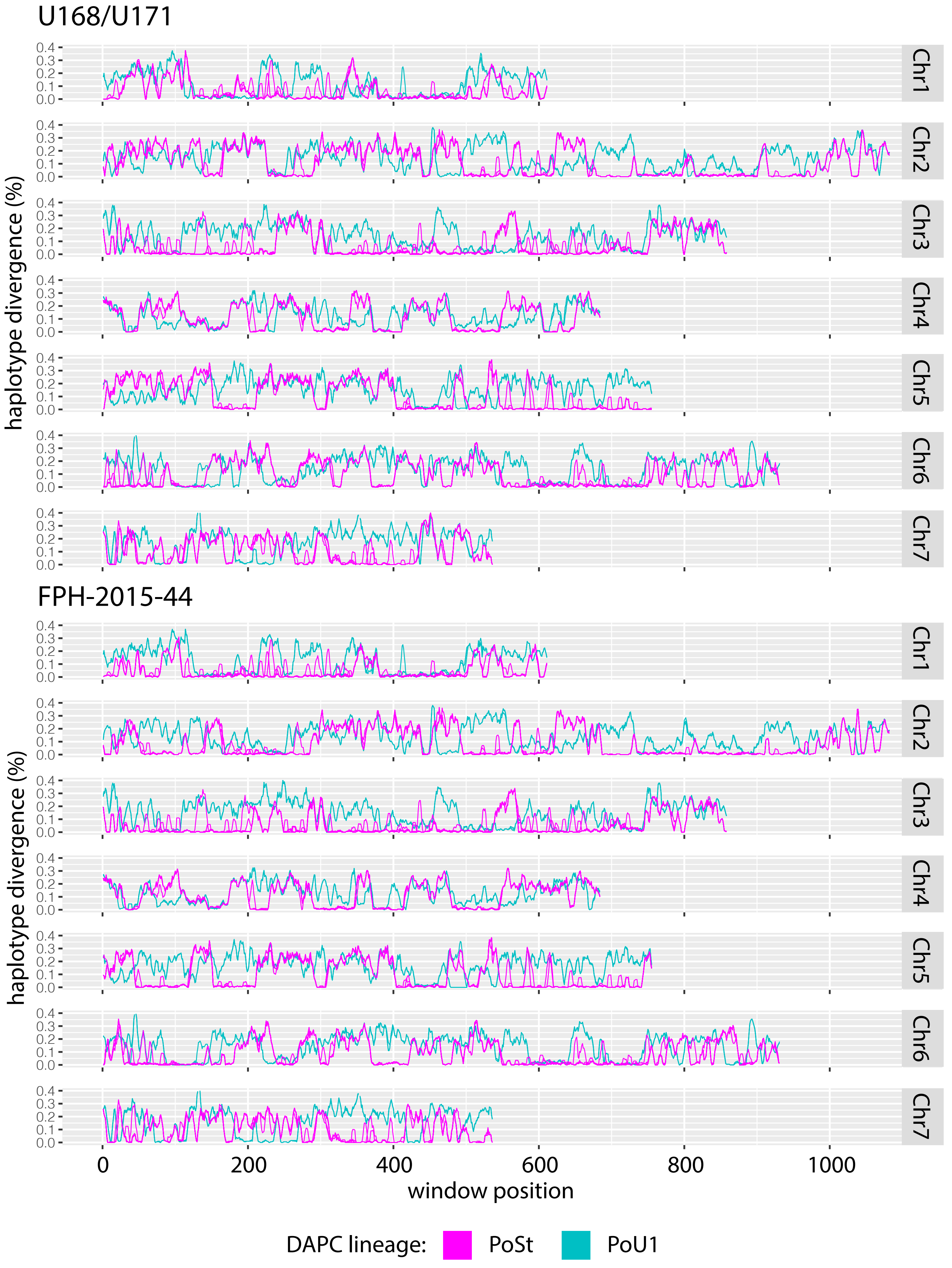
U168/U171
FPH-2015-44
haplotype divergence (%)
0.0
0.1
0.2
0.3
0.4
Chr1
Chr2
Chr3
Chr4
Chr5
Chr6
Chr7
0
200
400
600
800
1000
window position
DAPC lineage:
PoSt
PoU1

**Figure S5**

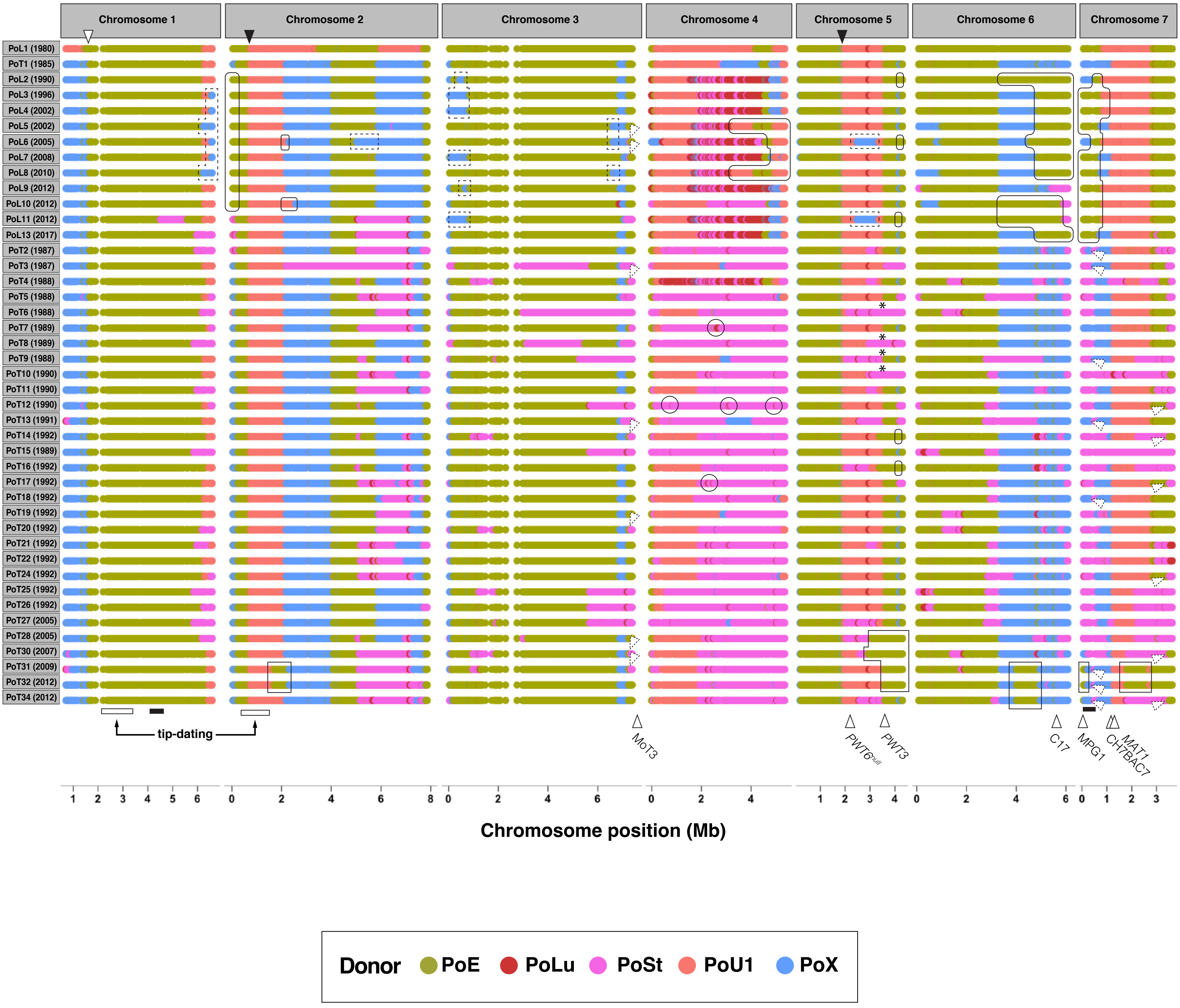

Chromosome 1
Chromosome 2
Chromosome 3
Chromosome 4
Chromosome 5
Chromosome 6
Chromosome 7
PoL1 (1980)
PoT1 (1985)
PoL2 (1990)
PoL3 (1996)
PoL4 (2002)
PoL5 (2002)
PoL6 (2005)
PoL7 (2008)
PoL8 (2010)
PoL9 (2012)
PoL10 (2012)
PoL11 (2012)
PoL13 (2017)
PoT2 (1987)
PoT3 (1987)
PoT4 (1988)
PoT5 (1988)
PoT6 (1988)
PoT7 (1989)
PoT8 (1989)
PoT9 (1988)
PoT10 (1990)
PoT11 (1990)
PoT12 (1990)
PoT13 (1991)
PoT14 (1992)
PoT15 (1989)
PoT16 (1992)
PoT17 (1992)
PoT18 (1992)
PoT19 (1992)
PoT20 (1992)
PoT21 (1992)
PoT22 (1992)
PoT24 (1992)
PoT25 (1992)
PoT26 (1992)
PoT27 (2005)
PoT28 (2005)
PoT30 (2007)
PoT31 (2009)
PoT32 (2012)
PoT34 (2012)
tip-dating
MoT3
PWT6null
PWT3
C17
MPG1
CH7BAC7
MAT1
Chromosome position (Mb)
Donor
PoE
PoLu
PoSt
PoU1
PoX

Figure S6

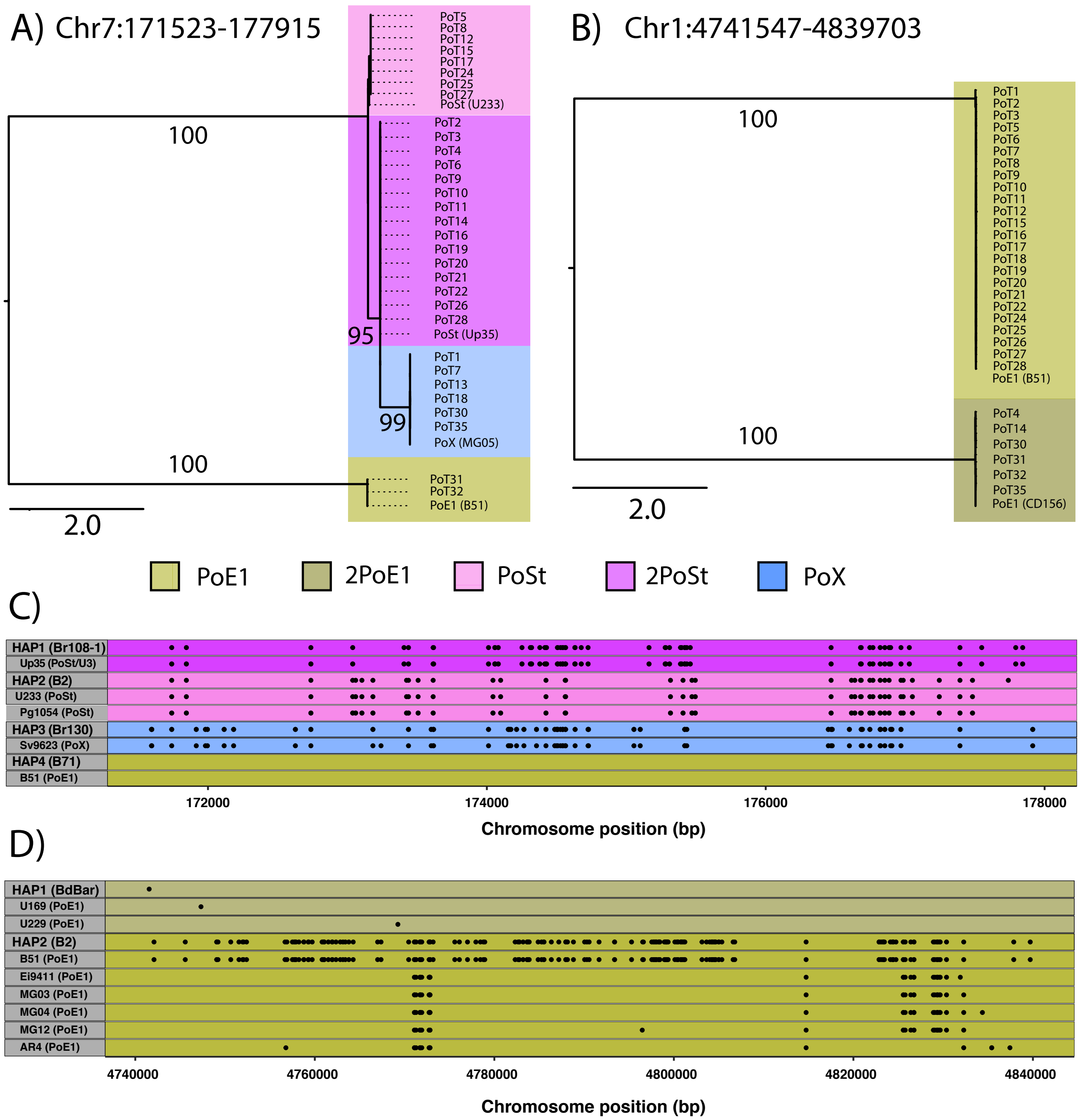

A) Chr7:171523-177915
B) Chr1:4741547-4839703
PoE1
2PoE1
PoSt
2PoSt
PoX
C)
D)
Chromosome position (bp)

**Figure S7**

A)

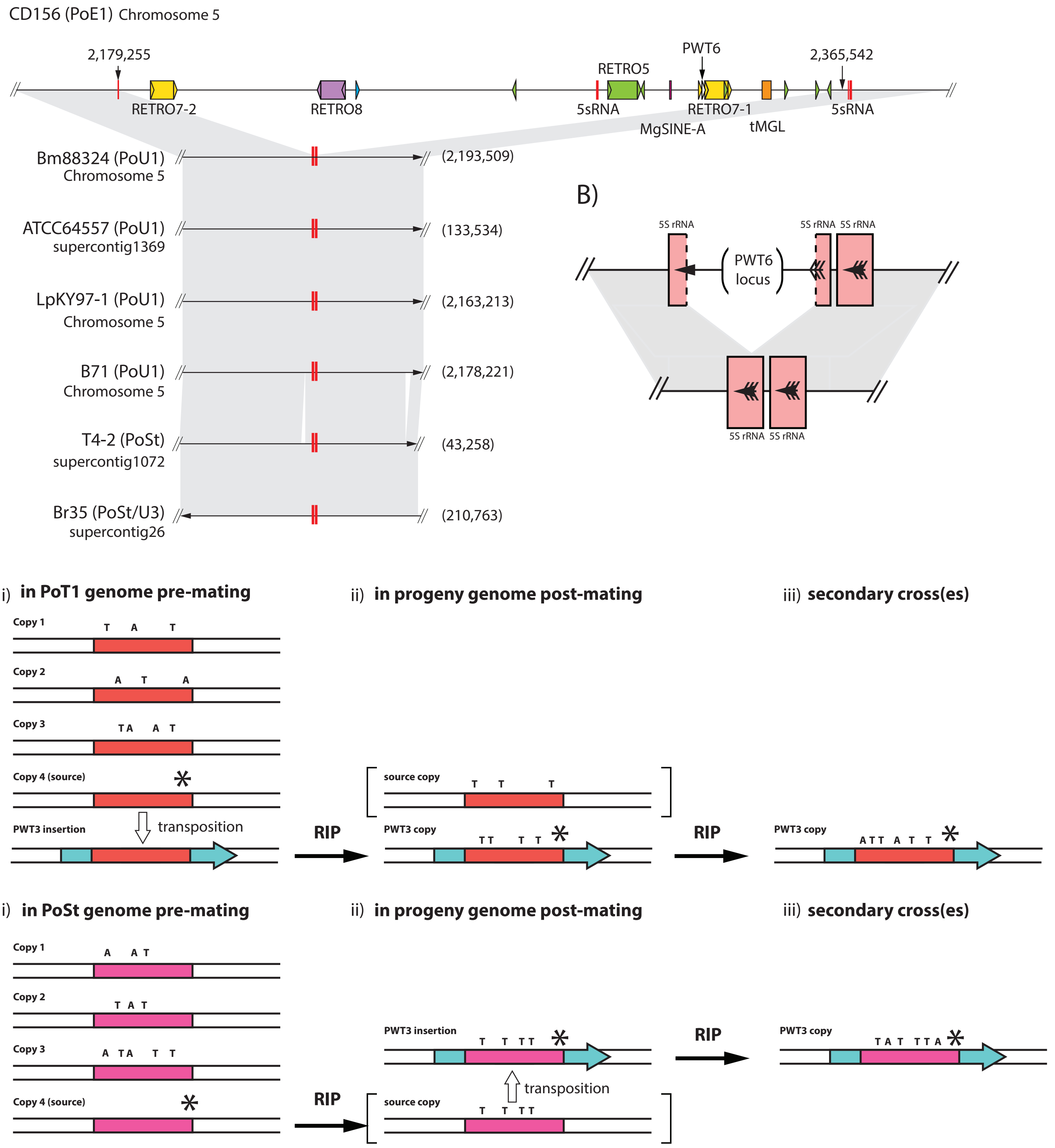

E)

| Query | Subject | qstart-qend | % id | sstart | send |
|---|---|---|---|---|---|
| Up35_contig491 | **Br81**_contig437 | 329492-335501 | 97.032 | 1 | 4448 |
| Up35_contig491 | **T12-8**_contig341 | 378958-384990 | 97.034 | 2175 | 5984 |

206CT88GA38GA50CT31GA139GA24CT15GA102CT4**TC**67GA60CT81CT8CT47CT89GA30GA23GA3GA4CT39GA1GA2CT9
GA8CT13GA9CT28GA3GA95GA23GA24GA34GA1GA36CT12CT26GA30CT12GA14GA31CT1CT6GACT10CT16GA14GA142

F)

| Query | Subject | qstart-qend | % id | sstart | send |
|---|---|---|---|---|---|
| T3-1_contig537 | **Br81**_contig437 | 329492-335501 | 95.551 | 1 | 1326 |
| T3-1_contig537 | **T12-8**_contig341 | 378958-384990 | 95.551 | 1 | 1326 |

5CT15GA83GA10**TC**7CT4**TC**40**AG**26GA60CT81CT8CT30**TC**16CT89GA54GA3GA4CT41GA2CT9GA8CT13GA9CT7**AGAG**19
GA3GA6TC13TC74GA23GA24GA36GA16**TC**19CT2**AG**34**TC**32CT12GA46CT1CT6GACT10CT16GA14GA52**AG**35AG44**AG**8
GA8GA18GA8CT14GACT20GA21GAGA6GA3

**Figure S8**
